# Actin assembly and non-muscle myosin activity drive dendrite retraction in an UNC-6/Netrin dependent self-avoidance response

**DOI:** 10.1101/492769

**Authors:** Lakshmi Sundararajan, Cody J. Smith, Joseph D. Watson, Bryan A. Millis, Matthew J. Tyska, David M. Miller

**Affiliations:** Department of Cell and Developmental Biology, Vanderbilt University, Nashville, TN; Department of Biological Sciences, Notre Dame University, South Bend, IN; Rho, Chapel Hill, NC; Neuroscience Graduate Program, Vanderbilt University, Nashville, TN; Cell Imaging Shared Resource, Vanderbilt University, Nashville, TN; Vanderbilt Biophotonics Center, Vanderbilt University, Nashville, TN

## Abstract

Dendrite growth is constrained by a self-avoidance response that induces retraction but the downstream pathways that balance these opposing mechanisms are unknown. We have proposed that the diffusible cue UNC-6(Netrin) is captured by UNC-40(DCC) for a short-range interaction with UNC-5 to trigger self-avoidance in the *C. elegans* PVD neuron. Here we report that the actin-polymerizing proteins UNC-34(Ena/VASP), WSP-1(WASP), UNC-73(Trio), MIG-10(Lamellipodin) and the Arp2/3 complex effect dendrite retraction in the self-avoidance response mediated by UNC-6(Netrin). The paradoxical idea that actin polymerization results in shorter rather than longer dendrites is explained by our finding that NMY-1 (non-muscle myosin II) is necessary for retraction and could therefore mediate this effect in a contractile mechanism. Our results also show that dendrite length is determined by the antagonistic effects on the actin cytoskeleton of separate sets of effectors for retraction mediated by UNC-6(Netrin) versus outgrowth promoted by the DMA-1 receptor. Thus, our findings suggest that the dendrite length depends on an intrinsic mechanism that balances distinct modes of actin assembly for growth versus retraction.

**AUTHOR SUMMARY:** Neurons may extend highly branched dendrites to detect input over a broad receptive field. The formation of actin filaments may drive dendrite elongation. The architecture of the dendritic arbor also depends on mechanisms that limit expansion. For example, sister dendrites from a single neuron usually do not overlap due to self-avoidance. Although cell surface proteins are known to mediate self-avoidance, the downstream pathways that drive dendrite retraction in this phenomenon are largely unknown. Studies of the highly branched PVD sensory neuron in *C. elegans* have suggested a model of self-avoidance in which the UNC-40/DCC receptor captures the diffusible cue UNC-6/Netrin at the tips of PVD dendrites where it interacts with the UNC-5 receptor on an opposing sister dendrite to induce retraction. Here we report genetic evidence that UNC-5-dependent retraction requires downstream actin polymerization. This finding evokes a paradox: How might actin polymerization drive both dendrite growth and retraction? We propose two answers: (1) Distinct sets of effectors are involved in actin assembly for growth vs retraction; (2) Non-muscle myosin interacts with a nascent actin assemblage to trigger retraction. Our results show that dendrite length depends on the balanced effects of specific molecular components that induce growth vs retraction.

## INTRODUCTION

Dendritic arbors are defined by the balanced effects of outgrowth which expands the structure versus retraction which constrains the size of the receptive field. Microtubules and filamentous actin (F-actin) are prominent dendritic components and have been implicated as key drivers of dendritic growth and maintenance by the finding that treatments that perturb cytoskeletal dynamics may also disrupt dendritic structure [1–3]. For many neurons, dendritic growth is highly exuberant with multiple tiers of branches projecting outward from the cell soma. Ultimately, growth may be terminated by external cues. For example, neurons with similar functions are typically limited to separate domains by tiling mechanisms in which mutual contact induces dendrite retraction. The related phenomenon of self-avoidance is widely employed to prevent overlaps among sister dendrites arising from a single neuron [4]. Homotypic interactions between the membrane components Dscam and protocaderins can mediate the self-avoidance response [5–7]. Surprisingly, in some instances, soluble axon guidance cues and their canonical receptors are also required. For example, self-avoidance for the highly-branched Purkinje neuron depends on both protocadherins and also repulsive interactions between sister dendrites decorated with the Robo receptor and its diffusible ligand, Slit [8]. In another example, we have shown that UNC-6/Netrin and its cognate receptors, UNC-5 and UNC-40/DCC, mediate self-avoidance for PVD nociceptive neurons in *C. elegans* [9]. Although multiple cell-surface interactions are now known to trigger self-avoidance, little is known of the downstream effectors that drive dendrite retraction in this mechanism.

The PVD neurons, one on each side of the body, build complex dendritic arbors through a series of successive 1°, 2°, 3° and 4° orthogonal branching events [10] [11–13]. Self-avoidance ensures that adjacent 3° branches do not overgrow one another [9,11] (Figure S1). Our previous results suggested a novel mechanism of self-avoidance in which UNC-40/DCC captures UNC-6/Netrin at the PVD cell surface and then triggers retraction by interacting with UNC-5 on the neighboring 3° branch. UNC-40/DCC may also effect self-avoidance by acting in a separate pathway that does not involve UNC-6/Netrin [9]. Here we describe a cell biological model of dendrite retraction in the PVD self-avoidance response that depends on actin polymerization.

The structure of the actin cytoskeleton is controlled by a wide array of effector proteins that regulate specific modes of actin polymerization. For example, Ena/VASP enhances F-actin elongation at the plus-end [12] and the Arp2/3 complex functions with WASP (Wiskott-Aldrich syndrome protein) and the Wave Regulatory Complex (WRC) to promote F-actin branching [13]. Upstream regulators of WASP and the WRC include Rho family GTPases [14] and their activators, the GEFs (Guanine nucleotide Exchange Factors) UNC-73/Trio [15,16] and TIAM (T-cell Lymphoma Invasion and Metastasis Factor) [17,18]. Members of the Lamellipodin/Lpd family recruit Ena/VASP to localize F-actin assembly at the leading edge of migrating cells [19]. The fact that all of these components (UNC-34/Ena/VASP, Arp2/3 complex, WSP-1/WASP, WRC, UNC-73/Trio, TIAM/TIAM-1, MIG-10/lamellipodin) have been previously shown to function in *C. elegans* to mediate axon guidance underscores the key role of the actin cytoskeleton in growth cone steering [15,18,20–23].

Our results show that UNC-34/Ena/VASP functions downstream of UNC-5 to mediate PVD self-avoidance. Genetic evidence detecting roles for the Arp2/3 complex and its upstream regulator, WASP/WSP-1, indicates that the formation of branched actin networks may contribute to dendrite retraction. A necessary role for actin polymerization is also indicated by the defective PVD self-avoidance response of mutants with disabled UNC-73/Trio or MIG-10/lamellipodin. We show that PVD self-avoidance also requires NMY-1/non-muscle myosin II which we propose effects dendrite retraction in a contractile mechanism that drives the reorganization of the nascent actin cytoskeleton. The necessary role for myosin could explain how actin polymerization can result in shorter rather than longer dendrites in the self-avoidance response. Thus, we propose that UNC-6/Netrin triggers self-avoidance by simultaneously stimulating actin assembly and non-muscle myosin activity in 3° dendrites.

Because dendrite growth depends on actin polymerization [24,25], our finding that the actin cytoskeleton is also necessary for dendrite self-avoidance points to different modes of actin polymerization for growth *vs* retraction. Recent work has shown that a multicomponent complex involving the PVD membrane proteins DMA-1 and HPO-30 drives dendritic growth by linking the PVD actin cytoskeleton to adhesive cues on the adjacent epidermis [26,27]. Dendrite elongation, in this case, depends on the WRC and TIAM-1, both of which are known to promote actin polymerization. In contrast, our work has revealed that effectors of actin polymerization that are required for 3° branch self-avoidance (e.g., UNC-34/Ena/VASP, MIG-10/lamellipodin, UNC-73/Trio, Arp2/3 complex, WSP-1/WASP) are not necessary for 3° dendrite outgrowth. Additionally, we show that UNC-6/Netrin signaling antagonizes the DMA-1-dependent mechanism of dendritic growth. These findings are significant because they suggest that overall dendrite length is defined by the relative strengths of opposing pathways that utilize separate sets of effectors to differentially regulate actin polymerization for either growth or retraction.

## RESULTS

### UNC-6/Netrin mediates PVD sister dendrite self-avoidance

To visualize PVD morphology, we utilized a previously characterized GFP marker driven by a PVD-specific promoter (*PVD*::GFP) (Figure S1A-B) [11]. Each PVD neuron adopts a stereotypical morphology characterized by a series of orthogonal junctions between adjacent branches: 1° dendrites extend laterally from the PVD cell soma; each 2° dendrite projects along the dorsal-ventral axis to generate a T-shaped junction comprised of two 3° dendrites each with either an anterior or posterior trajectory; terminal 4° dendrites occupy interstitial locations between the epidermis and body muscles. The net result of this branching pattern is a series of tree-like structures or menorahs distributed along the 1° dendrite. In contrast to the highly branched dendritic architecture, a single axon projects from the PVD cell soma to join the ventral nerve cord [10,11,26]. Despite the complexity of this network, dendrites rarely overlap (< 5%) in the mature PVD neuron (Figure S1A) [9]. Self-avoidance depends on an active process in which sister PVD dendrites retract upon physical contact with one another (Figure S1C). In the wild type, 3° branches arising from adjacent menorahs initially grow outward toward one another but then retract after mutual contact. The net result is a characteristic gap between the tips of 3° dendrites in neighboring menorahs [11]. We have previously shown that 3° dendrite self-avoidance depends in part on the diffusible cue UNC-6/Netrin and its canonical receptors, UNC-40/DCC and UNC-5 [9]. Mutations that disable *unc-6*, for example, result in a significant increase in the fraction of 3° dendrites that fail to retract and remain in contact (Figure S1B, Figure 3D). Similar self-avoidance defects were observed for mutants of either *unc-40* or *unc-5*. Our results are consistent with a model in which UNC-40 captures UNC-6 for contact with UNC-5 at the tips of adjacent 3° dendrites (Figure S1D)[9]. Here we address the question of how the activated UNC-5 receptor then modifies the actin cytoskeleton to drive dendrite retraction.

### Constitutively active UNC-5 delays dendrite outgrowth

We have proposed that UNC-6 initiates dendrite retraction by activating UNC-5 [9]. This model predicts that a constitutively active form of UNC-5 should retard net outgrowth of 3º dendrites by triggering an opposing retraction mechanism. To test this idea, we attached a myristoylation sequence to the cytoplasmic domain of UNC-5 [27]. The resultant myristyolated UNC-5 intracellular domain (MYR::UNC-5) is effectively tethered to the cytoplasmic membrane which is sufficient to activate UNC-5 dependent downstream signaling [27]. Expression of MYR::UNC-5 in PVD results in substantially shorter 3° dendrites in comparison to wild type at the L3 stage (Figure 1A-C). This phenotype suggests that constitutive activation of UNC-5 impairs normal outgrowth of 3° dendrites. We tested this idea by scoring the dynamic status of 3° dendrites. To quantify movement, we measured the distance from a fiduciary point to the tip of a 3° dendrite at intervals over time (delta t = 240 sec) (Figure 1E) (Movies S1-S2). Examples of these measurements are shown in Figure 1E. We observed that wild-type 3° dendrites typically display saltatory movement with periodic bouts of extension and retraction that result in net elongation. In contrast, 3° dendrites of PVD neurons expressing MYR::UNC-5 grow more slowly and rarely show periods of active extension and retraction (Figures 1E and 1F). The higher overall motility of 3° dendrites in the wild-type vs PVD::MYR::UNC-5 strain is evident from a comparison of the standard deviation of the net movement of 3° dendrites from a fiduciary point in each strain (Figure 1F). Despite slower outgrowth, 3° dendrites expressing MYR::UNC-5 eventually produce a normal-appearing PVD architecture by the late L4 stage (Figure 2A). MYR::UNC-5 lacks the extracellular UNC-6-binding domain (Figure 1D) and thus should function independently of UNC-6. We confirmed this prediction by demonstrating that MYR::UNC-5 prevents the self-avoidance defect observed in an *unc-6* mutant in L4 stage animals (Figure 2B, D). We attribute this effect to the constitutive activation of dendrite retraction by MYR::UNC-5 that effectively prevents overgrowth of adjacent sister 3° dendrites in an *unc-6/Netrin* mutant. Together, these results are consistent with the hypothesis that UNC-5 functions downstream of UNC-6/Netrin to trigger dendrite retraction in the self-avoidance response.

**Figure 1:**
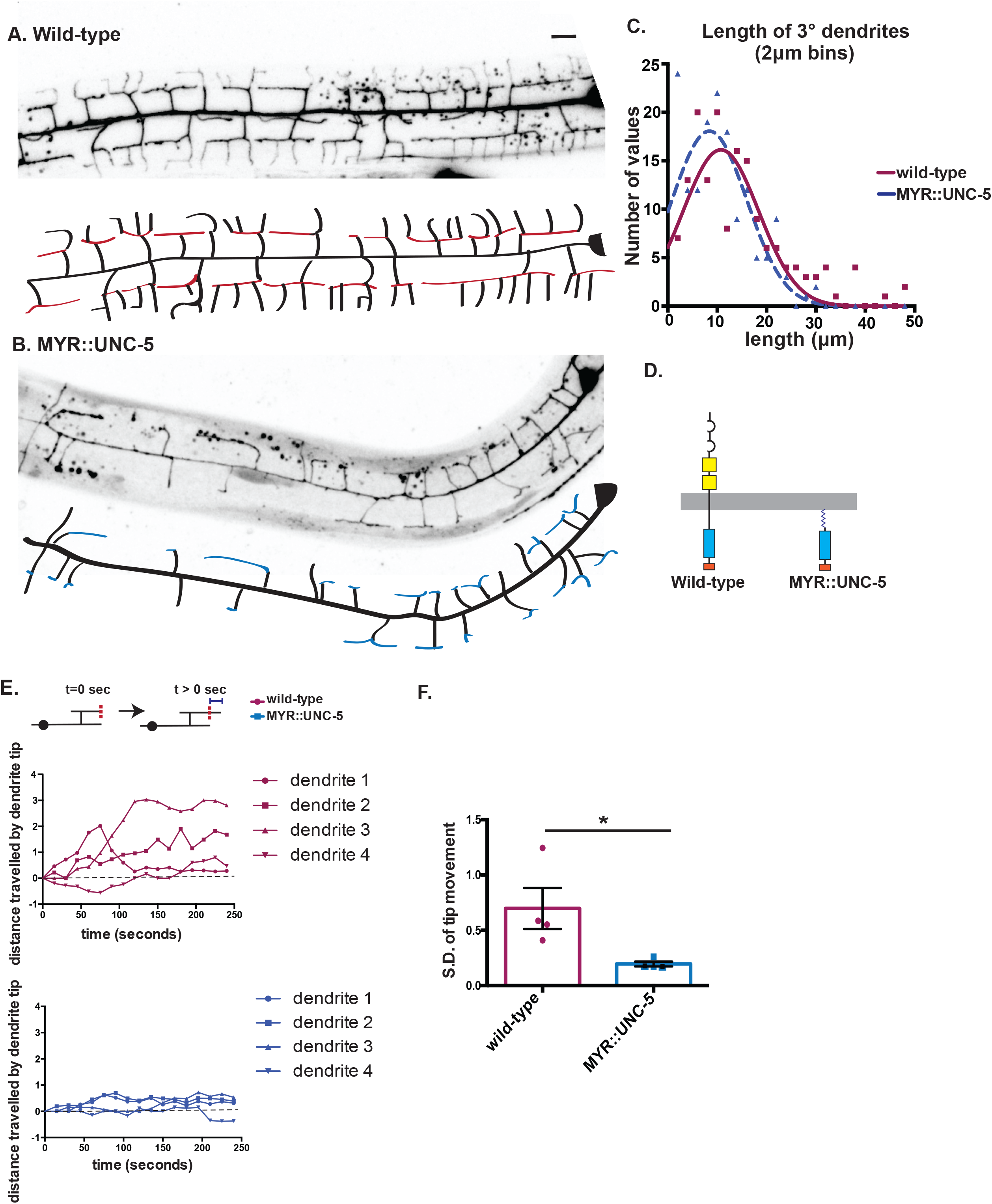
Constitutively active UNC-5 retards outgrowth of 3° PVD dendrites. (A-D) Representative images and tracings of 3° dendrites anterior to PVD cell body in (A) wild type (red) and (B) PVD::MYR::UNC-5 (blue) (Late L3-early L4 animals) (C) Gaussian fit to distribution of lengths of PVD::mCherry-labeled 3° dendrites showing shorter average length for MYR::UNC-5 (n = 156, 7 animals) *versus* wild type (n = 159, 8 animals), p = 0.03, Kolmogorov-Smirnov statistical test. (D) Schematics of intact, wild-type UNC-5 receptor vs membrane-associated, constitutively-activated myristylated-UNC-5 (MYR::MYR-5). (E-F) Dynamicity of 3° dendrites in wild-type vs MYR::UNC-5-expressing PVD neurons labeled with PVD::GFP. (E) Schematic (top) depicts displacement of the tip of a 3° dendrite relative to a fiduciary mark (vertical dashed red line). Scale bar is 10 μm. Representative plots of displacement for wild-type (red) and MYR::UNC-5 (blue) 3° dendrites over time. (F) The dynamic saltatory growth and retraction of 3° dendrites in the wild type vs the relative quiescence of MYR::UNC-5-expressing PVD neurons is correlated with elevated standard deviation of tip movement from a fiduciary point in wild-type vs MYR::UNC-5 expressing PVD neurons (p = 0.04 from Student’s t-test. n = 4).

**Figure 2:**
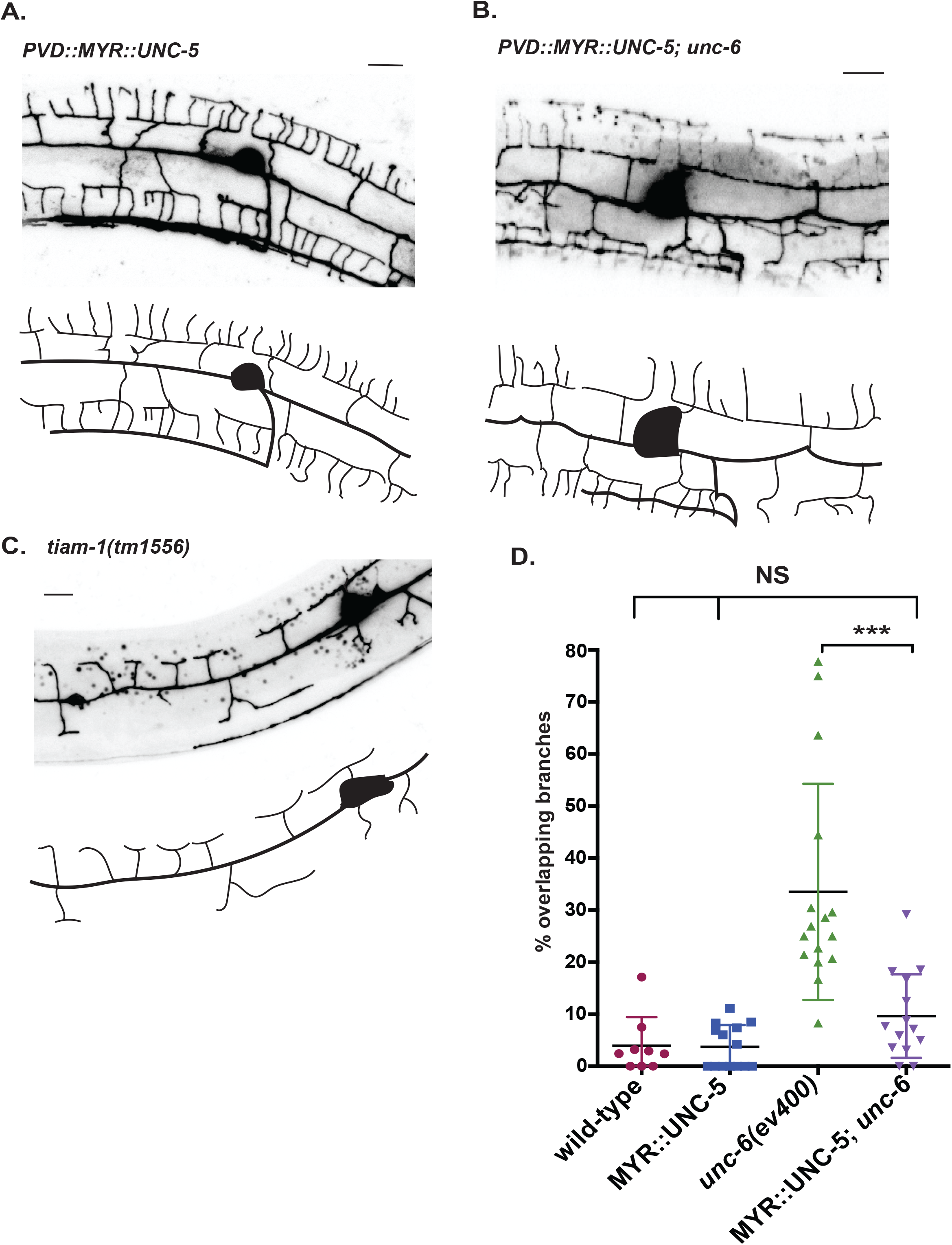
Constitutively active UNC-5 rescues self-avoidance in an *unc-6* mutant. (A-C) Representative images (top) and tracings (bottom) of PVD dendritic architecture in late L4 stage larvae that are (A) PVD::MYR:UNC-5, (B) PVD::MYR::UNC-5; *unc-6(ev400)*, (C) Self-avoidance defects (percentage of overlapping 3° branches) are rare (< 5%) in wild type and PVD::MYR::UNC-5. The Unc-6 self-avoidance defect is prevented by PVD expression of MYR::UNC-5. Error bars denote mean and M. ***p<0.0001. n > 15. from ANOVA with Tukey’s posthoc test. Scored in a PVD::mCherry background.

### UNC-34/Ena/VASP mediates self-avoidance in actin-containing PVD dendrites

To explore potential modifications of the PVD cytoskeleton as effectors of the UNC-6/Netrin-dependent self-avoidance response, we used live cell markers to visualize actin (PVD::ACT-1::GFP) and microtubules (*pdes-2*::TBA-1::mCherry) in PVD [28]. We observed a robust ACT-1::GFP signal throughout the PVD neuron including the axon and all dendrites (Figure 3A). In contrast, the TBA-1::mCherry signal was strongest in the PVD axon and 1° dendrite. This result confirms an earlier report suggesting that dendritic microtubules are most abundant in the 1° branch (Figure 3A) [28]. We thus considered the possibility that the self-avoidance response could depend on the regulation of actin dynamics. To test this idea, we examined the PVD self-avoidance phenotype for a mutation that disables the actin binding protein, Ena/VASP. We selected Ena/VASP for this test because it has been shown to function as a cytoplasmic effector of actin polymerization downstream of *unc-6/Netrin* signaling [12,29,30]. In addition, the *C. elegans* Ena/VASP homolog, UNC-34 [31], is enriched in a gene expression profile of PVD neurons [11]. A strong loss-of-function allele of *unc-34*/Ena/VASP [32] alters overall PVD morphology (e.g., truncated or misplaced 1°dendrite, fewer 2° branches) (Figure 3B-C, Figure S2A-C). In contrast, mutations in other regulators of actin polymerization (e.g. TIAM-1/Gef, Figure S2D) result in severe truncation of lateral branches including 3° dendrites, a phenotype that points to a role for actin polymerization in dendrite outgrowth [24,25]. PVD neurons in *unc-34* mutant animals show the opposite effect of adjacent 3° dendrites that fail to self-avoid and overgrow one another (Figure 3B-C). PVD-specific expression of mCherry-tagged UNC-34 protein rescued this Unc-34 mutant phenotype suggesting that *unc-34* function is cell-autonomous for the self-avoidance response (Figure 3C). *unc-34* mutant 1°and 2° branch defects were not rescued by PVD expression of UNC-34, however, which could be indicative of an additional UNC-34 function in nearby cell types that influence PVD morphogenesis (Figure 3B, Figure S2) [33,34]. Thus, our results suggest that UNC-34/Ena/VASP is selectively required in PVD neurons for dendrite retraction in the self-avoidance response but not for dendrite outgrowth. The previously established role of UNC-34/Ena/VASP as a downstream effector of UNC-5 [20] predicts that *unc-34* should be epistatic to the MYR::UNC-5 gain-of-function phenotype. We confirmed this prediction by showing that the delayed outgrowth PVD dendrites in the PVD::MYR::UNC-5 strain is fully suppressed in *unc-34* mutant backgrounds which display the Unc-34 self-avoidance defect (Figure 3C). This result stands in striking contrast to our finding above that PVD::MYR::UNC-5 is epistatic to the Unc-6 self-avoidance phenotype (Figure 2B, C). Thus, our genetic results are consistent with a model in which UNC-6/Netrin acts through UNC-5 to activate UNC-34/Ena/VASP for dendrite retraction in the self-avoidance response.

**Figure 3.**
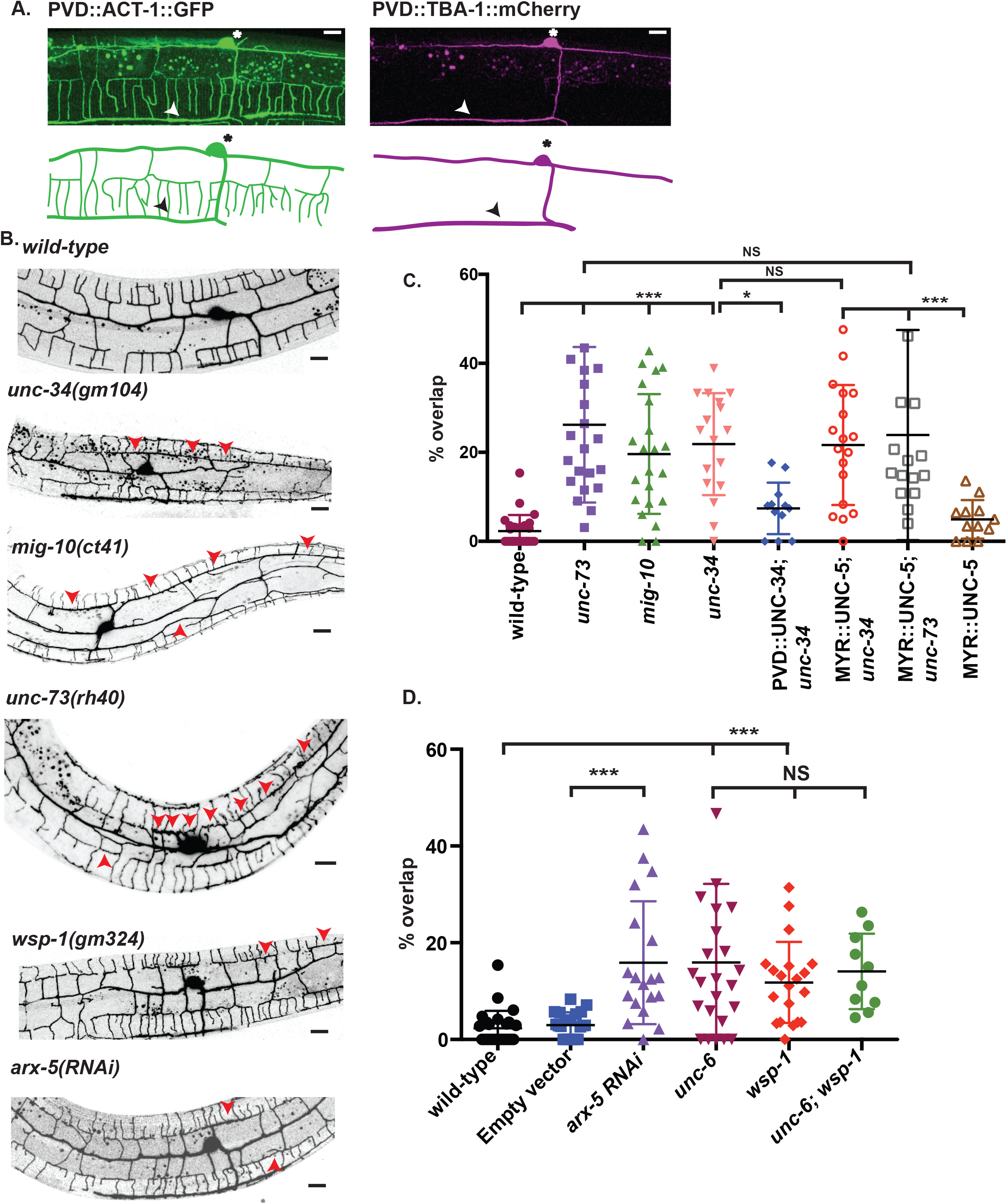
Mutations in genes that promote actin polymerization disrupt PVD dendrite self-avoidance. (A) Representative images (top) and schematics (bottom) of the actin cytoskeleton (green) labeled with PVD::ACT-1::GFP (A) and microtubules (magenta) marked with *pdes*-*2*::TBA-1::mCherry. PVD cell bodies (asterisks) and axons (arrowheads) are noted. (B) Representative images of PVD::GFP for wild type, for *unc-34*, *mig-10*, *unc-73*, *wsp-1* mutants and for *arx-5RNAi*. Asterisks denote PVD cell bodies and arrowheads point to overlaps between adjacent PVD 3° dendrites. (C-E) Self-avoidance defects are plotted in a PVD::GFP background (mean and SEM) as the percentage of overlapping 3° dendrites for each genotype. ***p<0.001, *p<0.05, ANOVA with Tukey’s posthoc test, n > 15 animals for all backgrounds. (C) Self-avoidance defects for *unc-73/Trio*, *mig-10/Lmpd* and *unc-34/Ena/VASP* mutants. Self-avoidance defects of *unc-34/Ena/VASP* mutants are rescued by PVD expression of UNC-34 (PVD::mCherry::UNC-34). Constitutively active UNC-5 (PVD::MYR::UNC-5) fails to rescue the self-avoidance defects of an *unc-34* mutant. (D) PVD self-avoidance defects for *arx-5RNAi* compared to Empty Vector RNAi controls., ***p<0.001 and self-avoidance defect of *unc-6*, *wsp-1* and *unc-6; wsp-1* mutants compared to wildtype, NS = Not Significant, ANOVA with Tukey’s posthoc test, n > 15 animals. Scale bar is 10 μm.

For visualizing UNC-34/Ena/VASP in developing dendrites, we labeled UNC-34 with mCherry and co-expressed it in PVD (PVD::mCherry::UNC-34) with LifeAct::GFP to detect actin-containing structures. mCherry::UNC-34 is functional because it rescues the *unc-34* self-avoidance defect (Fig 3C). iSIM super resolution imaging detected punctate mCherry::UNC-34 throughout the PVD dendritic array (see Methods) Additional images were collected by AiryScan and TIRF microscopy (Figure S3). Notably, mCherry::UNC-34 puncta are consistently detected at the tips of 3° dendrites (Figure 4A-C, S3), a finding consistent with the observed distal location of Ena/VASP in F-actin containing filopodial structures [35,36]. We used time-lapse imaging to detect potential trafficking of mCherry::UNC-34 (Movie S5). Kymographs as well as particle tracking analysis, detected both mobile and stationary mCherry::UNC-34 puncta in 3° dendrites (Figure 4D-F, S3). These results are consistent with the idea that UNC-34/Ena/VASP is delivered to the tips of 3° dendrites where it mediates actin assembly for branch retraction in the self-avoidance response

**Figure 4.**
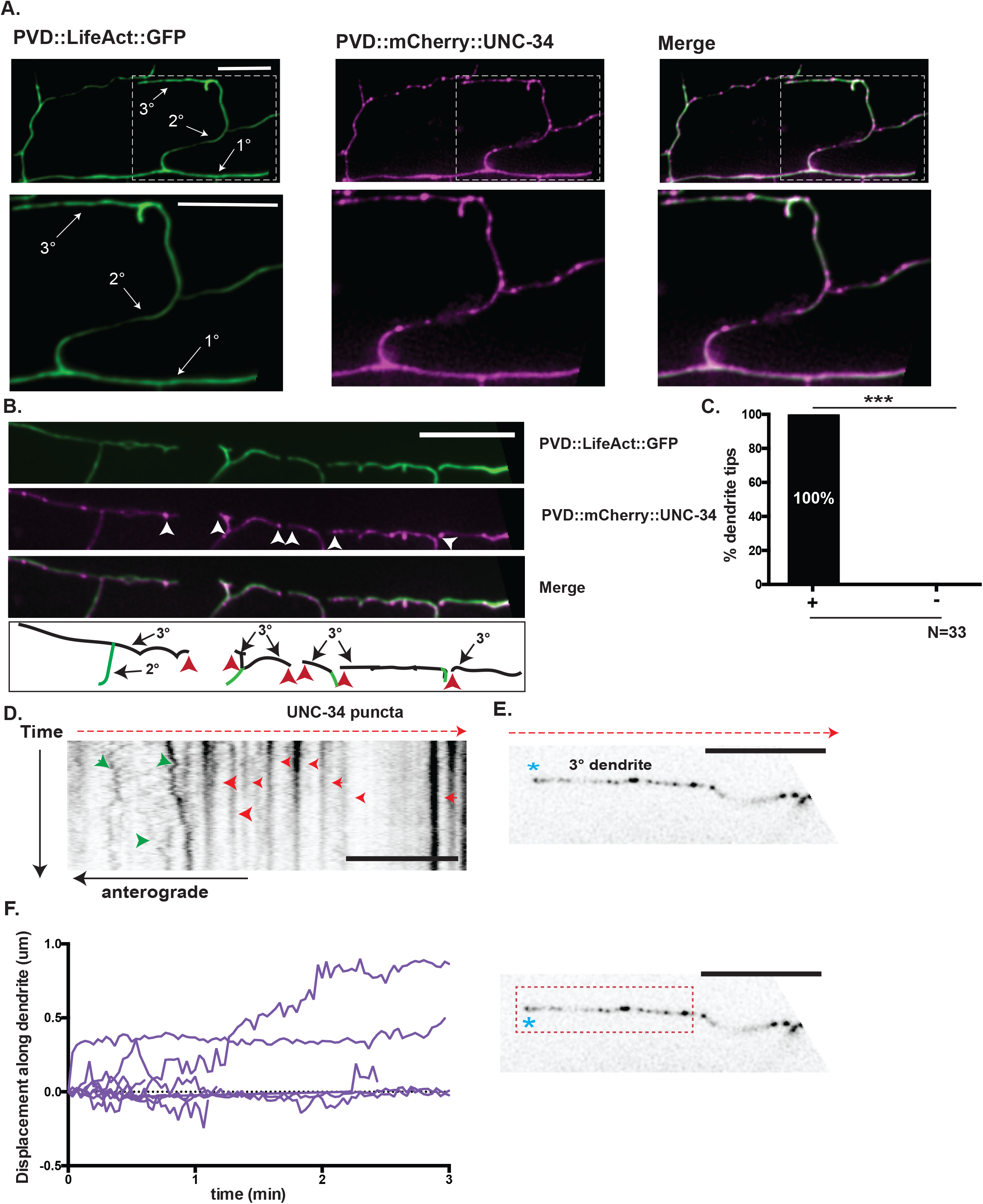
Localization and trafficking of PVD::UNC-34::mCherry puncta in PVD. (A-B) iSIM images (see Methods) of PVD neurons (late L3 larvae) labeled with PVD::LifeAct::GFP (green) and PVD::mCherry::UNC-34 (magenta). Merge shown at top right. Insets depict punctate mCherry::UNC-34 in 1°, 2° and 3° PVD dendrites. (B) Note mCherry::UNC-34 puncta at tips (white arrowheads) of 3° dendrites. Scale bars are 5 μm. (C) Frequency (100%) at which tips of 3º dendrites show mCherry::UNC-34 puncta, ***p<0.001, Fischer’s exact test, N = 33 dendrite tips in 9 animals. Both iSIM and Airyscan images were used in this analysis. (D-E) Kymographs of PVD::mCherry::UNC-34 in PVD 3° dendrite captured in the iSIM microscope. Moving (green arrowheads) vs stationary (red arrowheads) mCherry::UNC-34 puncta are denoted. The distal end of the 3° dendrite is indicated by an asterisk and a red dashed arrow points toward the PVD cell soma. (F) Trafficking of mCherry::UNC-34 puncta in the PVD 3° dendrite (enclosed with rectangle of red dashed line) from 4E was analyzed using a spots tracking algorithm (Imaris) (see Methods). The net displacement (µm) of mobile mCh::UNC-34 puncta (N = 8) from position at t = 0 in a 3º dendrite plotted against time with a sampling interval of 2 s. Scale bar is 10 μm.

### Mutations that disable actin-polymerizing proteins disrupt dendrite self-avoidance

Genetic tests of additional regulators of actin polymerization also revealed PVD self-avoidance phenotypes. Lamellipodin (Lpd) interacts with ENA/VASP at the tips of filopodia and lamellopodia to extend actin filaments [12,19]. Mutants of *mig-10*/Lpd display robust PVD self-avoidance defects (Figure 3B, E). Similarly, self-avoidance is disrupted by a mutation that eliminates the Rac GEF activity of UNC-73/Trio, a potent regulator of actin dynamics [15]. The predicted downstream function of UNC-73/Trio in the UNC-5 signaling pathway [27] is confirmed by our finding that the self-avoidance defects of *unc-73/Trio*, like *unc-34/Ena/VASP* mutants, are fully epistatic to the delayed outgrowth phenotype of constitutively active PVD::MYR::UNC-5 (Figure 3). We note that both *mig-10*/Lpd and *unc-73/*Trio mutants also show other defects in PVD morphology (e.g., displaced 1° dendrite, fewer 2° branches) (Figure S2) closely resembling those of *unc-34*/Ena/VASP mutants which our results suggest are due to a non-cell autonomous function for UNC-34 (Figure S2).

To test for potential roles for branched actin polymerization, we examined a null allele of *wsp-1/WASP* (Wiskott-Aldrich Syndrome protein) and observed defective PVD self-avoidance. If *wsp-1*/WASP functions in a common pathway with *unc-6*, then the self-avoidance defect of the double mutant, *wsp-1; unc-6*, should be no more severe than that of either *wsp-1* or *unc-6* single mutants. We confirmed this prediction by scoring PVD self-avoidance in *wsp-1; unc-6* mutant animals (Figure 3D). An independent study confirmed the role of *wsp-1* in PVD self-avoidance. This work also reported, however, that *wsp-1* does not enhance the self-avoidance defect of a parallel-acting pathway involving MIG-14/Wntless and thus concluded that *wsp-1* functions downstream of MIG-14/Wntless [37]. Additional experiments are needed to resolve the question of whether WSP-1/WASP functions exclusively in the UNC-6/Netrin vs MIG-14/Wntless self-avoidance pathways.

Because WASP is known to activate the Arp2/3 complex [14,38], we also performed RNAi knock down of the conserved Arp2/3 component ARX-5/p21 and detected significant PVD self-avoidance defects (Figure 3E). The canonical roles of WASP and the Arp2/3 complex suggest that self-avoidance depends on the formation of branched F-actin networks. Recent studies have shown that actin and actin regulators like TIAM-1/GEF and components of the Wave Regulator Complex (WRC) are required for PVD dendrite outgrowth [26,27] (Figure S2D). Here our genetic results point to the paradoxical idea that elongation and branching of F-actin filaments are also necessary for dendrite retraction and that this mechanism is activated by UNC-6/Netrin in the self-avoidance response.

### The actin cytoskeleton is highly dynamic in growing and retracting PVD dendrites

To test the idea that the actin cytoskeleton is differentially modulated for both outgrowth and retraction we expressed LifeAct::GFP in PVD neurons to monitor actin dynamics during dendrite morphogenesis. Although LifeAct::GFP binds both filamentous actin (F-actin) and monomeric G-actin with high affinity, bright LifeAct::GFP-labeled foci typically correspond to F-actin-containing structures [39]. We observed that the LifeAct::GFP signal was consistently brightest nearest the tips of growing dendrites in comparison to internal dendritic regions (Fig. 5B, C) as also reported in other studies [26,27,40]. Time-lapse imaging of PVD::LifeAct::GFP by spinning disk confocal microscopy during the late L3/early L4 stage detected striking, dynamic fluctuations in the LifeAct::GFP signal (Movie S3). We quantified this effect by comparing time-dependent changes in LifeAct::GFP fluorescence versus that of a PVD::mCherry cytosolic marker. The relatively constant intensity of PVD::mCherry signal in these experiments favors the idea that the observed changes in LifeAct::GFP fluorescence could be due to actin dynamics in growing dendrites (Figure 5A, D–E).

**Figure 5.**
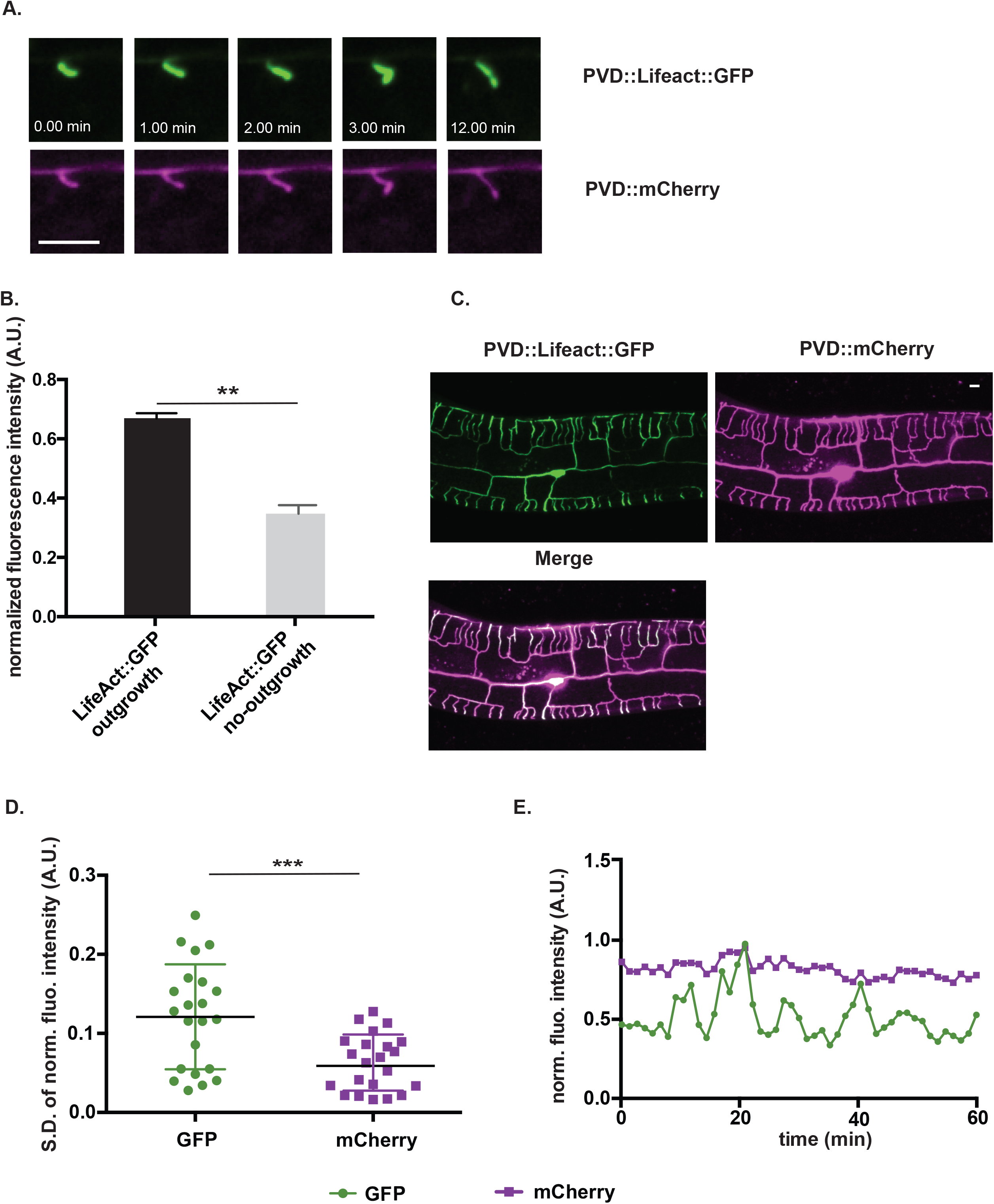
The actin cytoskeleton is highly dynamic in PVD dendrites. (A) Time-lapse images of PVD::LifeAct::GFP and cytosolic PVD::mCherry labeled 2° dendrite during outgrowth (late L3 larvae). (B) Fluorescent intensity was measured from regions of PVD dendrites undergoing outgrowth vs no-outgrowth (see Materials and Methods). ***p<0.001, N = 118 measurements from 6 dendrites. (C) Representative image showing enriched LifeAct::GFP signal in distal, growing PVD dendrites labeled with PVD::mCherry. (D) Rapid changes in the normalized fluorescence intensity of PVD::LifeAct::GFP versus the PVD::mCherry cytoplasmic marker were quantified as the standard deviation of the normalized fluorescent intensity measured at the tips of growing dendrites. (See Movie S3 and Methods) ***p = 0.003, N = 22. (Unpaired t-test) (E) Normalized fluorescence intensity measurements of PVD::LifeAct::GFP (green) vs cytoplasmic PVD::mCherry (red) at the tip of a growing late L3 stage PVD 3°dendrite from a representative time lapse movie.

Because genetic knockdown of actin polymerizing proteins (UNC-34/Ena/VASP, MIG-10/Lpd, UNC-73/Trio, WSP-1/WASP, ARX-5/P21) (Figure 3) disrupts self-avoidance and thus suggests that F-actin is required, we monitored the LifeACT::GFP signal at the tips of 3° dendrites during the self-avoidance response (Figure S4) (Movie S3). The cytoplasmic mCherry marker was used to detect contact events as instances in which the measured gap between left and right 3° dendrites approached zero. Although time-lapse imaging detected dynamic fluctuations in LifeAct::GFP fluorescence both before and after contact, we did not observe a quantitative increase in the LifeACT::GFP signal with retraction (Figure S4 and S5). Notably, for these experiments, we used the marker strains (i.e., PVD::LifeAct::GFP and PVD::mcherry), imaging strategy (i.e., 40 X objective, 35 s sampling interval) and method of quantitative analysis described in an earlier publication that reported increased LifeAct::GFP fluorescence after contact (N = 14, p = 0.54, two-way ANOVA with Bonferroni correction [37] (Figure S4A-C). In an additional approach, we used a higher resolution objective (100X) and longer sampling interval (1 – 1.3 min) and also did not detect elevated LifeAct::GFP fluorescence during retraction (N = 8, p = 0.74, two-way ANOVA with Bonferroni correction) (Figure S4D and S5). The simplest explanation of our results is that actin polymerization drives both outgrowth and retraction which are tightly coupled over time at dendrite tips and thus difficult to resolve as separate events.

### NMY-1/Non-muscle myosin II drives dendrite retraction in the self-avoidance mechanism

Our imaging results indicate that F-actin is the dominant cytoskeletal structure in growing PVD dendrites. In considering a mechanism to account for dendrite shortening in the self-avoidance response, we hypothesized that a newly synthesized actin cytoskeleton could mediate dendrite shortening by providing a substrate for non-muscle myosin II to drive contraction. To test this idea, we examined loss-of-function alleles that disable the *C. elegans* non-muscle myosins, NMY-1 and NMY-2. This experiment determined that mutations in *nmy-1* but not *nmy-*2 [40] show PVD self-avoidance defects (Figure 6). This effect is cell autonomous for *nmy-1* because PVD-specific RNAi of *nmy-1* also results in overlapping 3° dendrites and PVD-specific expression of GFP-tagged NMY-1 is sufficient to rescue the *nmy-1* self-avoidance defect (Figure 6 B–D). We conclude that a specific non-muscle myosin, NMY-1, promotes dendrite retraction in the self-avoidance mechanism and could act by driving the translocation of F-actin bundles at the tips of 3° dendrites.

**Figure 6.**
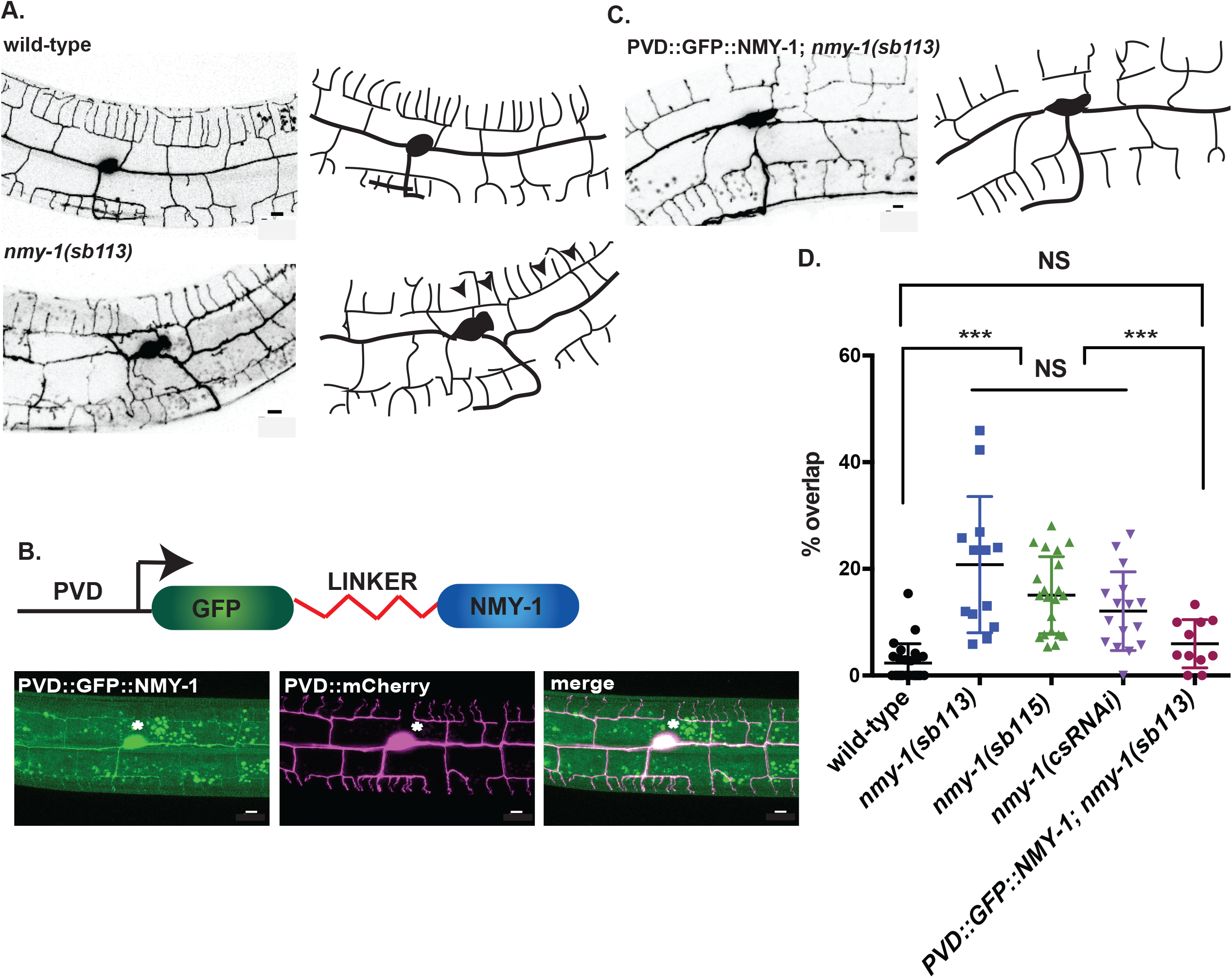
NMY-1/non-muscle myosin II mediates dendrite self-avoidance. (A) Representative images (left) and tracings (right) of wild type and *nmy-1(sb113)* labeled with PVD::GFP. Arrowheads mark overlapping 3° dendrites. The PDE neuron (asterisk) is adjacent to the PVD cell-body. (B) NMY-1 fused to GFP via a linker peptide (PVD::GFP::NMY-1) (top) expressed in PVD with PVD::mCherry (bottom) rescues self-avoidance defects (C) of *nmy-1(sb113).* Scale bars are 5μm. (D) Quantification of self-avoidance defects in wild type and *nmy-1*. The percentage of overlapping 3° branches is elevated in *nmy-1* mutants and with *nmy-1* cell-specific RNAi (csRNAi) vs wild type. PVD expression of GFP-labeled NMY-1 (PVD::GFP::NMY-1) rescues the *nmy-1(sb113)* self-avoidance defect. Scatter plots with mean and SEM, ***p<0.01, One way ANOVA with Tukey’s posthoc test. N = 15-20 animals per genotype. All defects were quantified in a PVD::GFP background. Scale bar is 5 μm.

### Phosphomimetic activation of MLC-4 shortens PVD 3° dendrites

Non-muscle myosin II motor proteins are composed of myosin heavy chain (MHC) and light chains, the essential light chain (ELC) and regulatory light chain (RLC) [41,42]. Myosin motor activity is triggered by RLC phosphorylation that induces a conformational change to allow assembly of the myosin complex into active bipolar filaments (Figure 7A) [43]. Having shown that NMY-1/non-muscle myosin is required for self-avoidance, we next performed an additional experiment to ask if constitutive non-muscle myosin activity is sufficient to induce dendrite retraction. For this test, we expressed a phosphomimetic mutant of MLC-4, the *C. elegans* RLC homolog. MLC-4/myosin regulatory light chain contains serine/S and threonine/T phosphorylation sites in a highly-conserved domain. Both S and T residues were mutated to Aspartate/D and the resultant phosphomimetic construct MLC-4DD was fused to GFP for transgenic expression in PVD (Figure 7B-C) [44]. PVD neurons that expressed MLC-4DD showed significantly shorter 3° dendrites at the L3 stage in comparison to wild type as well as a concurrent increase in the width of gaps between 3° dendrites from adjacent menorahs (Figure 7D-G). These phenotypic traits are consistent with the idea that normal dendrite outgrowth is slowed by constitutive activation of non-muscle myosin by MLC-4ADD. By the L4 stage, PVD architecture in the MLC-4DD strain resembles that of the wild type with normal appearing menorah morphology and spacing (Figure 8A, B). Similar results were observed for PVD neurons that express constitutively active MYR::UNC-5 which we have proposed functions downstream of UNC-6/Netrin to trigger the self-avoidance response (Figure 2). Our finding that MLC-4DD rescues the self-avoidance defect of *unc-6* mutants suggests that NMY-1/non-muscle myosin II also functions downstream of *unc-6* for dendrite retraction (Figure 8C).

**Figure 7.**
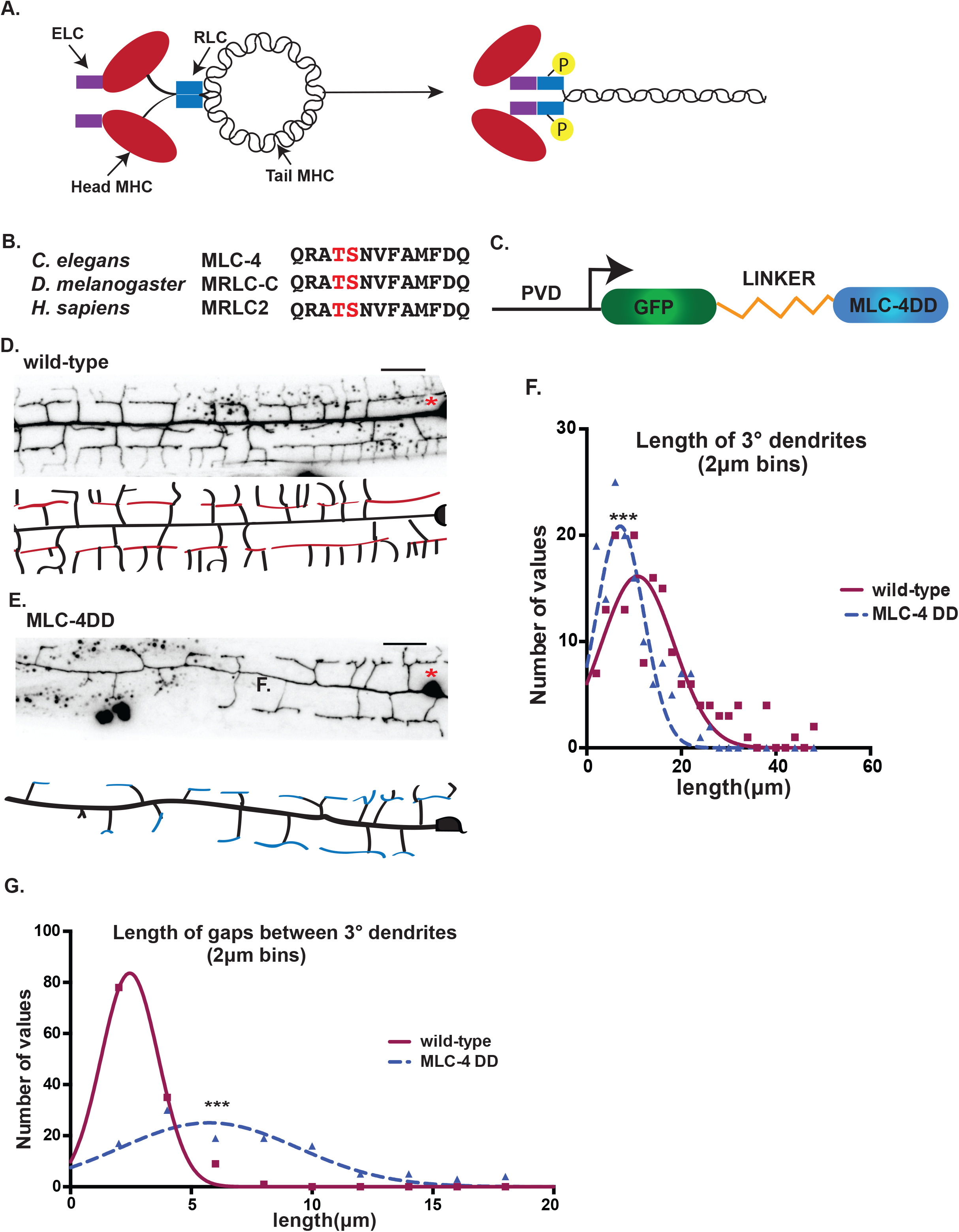
Phosphomimetic activation of myosin regulatory light chain, MLC-4, retards outgrowth of 3° PVD dendrites. (A) Phosphorylation of Regulatory Light Chain (RLC) induces a conformational change that activates the myosin protein complex composed of myosin heavy chain (MHC), essential light chain (ELC) and RLC. (B) Threonine and serine phosphorylation sites (red) in conserved regions of *C. elegans*, *Drosophila* and human myosin RLCs. (C) Phosphomimetic construct, PVD::GFP::MLC-4-DD, fused to GFP with a linker peptide for expression with PVD promoter. (D-E) Representative images (top) and schematics (bottom) of PVD morphology anterior to cell body (asterisk) denoting 3° dendrites for (D) wild type (red) and (E) MLC-4DD (blue) (L3 stage larvae). Wild-type image also used in Figure 2A. Scale bars are 10 μm. 3° dendrite length (F) is shorter and gaps between 3° dendrites (G) is wider for MLC-4DD versus wild type at the L3 stage. Gaussian fit was plotted with bin center values (μm) on the X-axis and the number of values that fall under each bin center on the Y axis, ***p<0.001, Kolmogorov-Smirnov test for cumulative distributions of PVD 3° dendrite lengths and length of gaps in MLC-4DD (n = 141, 6 animals) versus wild type (n = 156, 8 animals). Scale bar is 10 μm.

**Figure 8.**
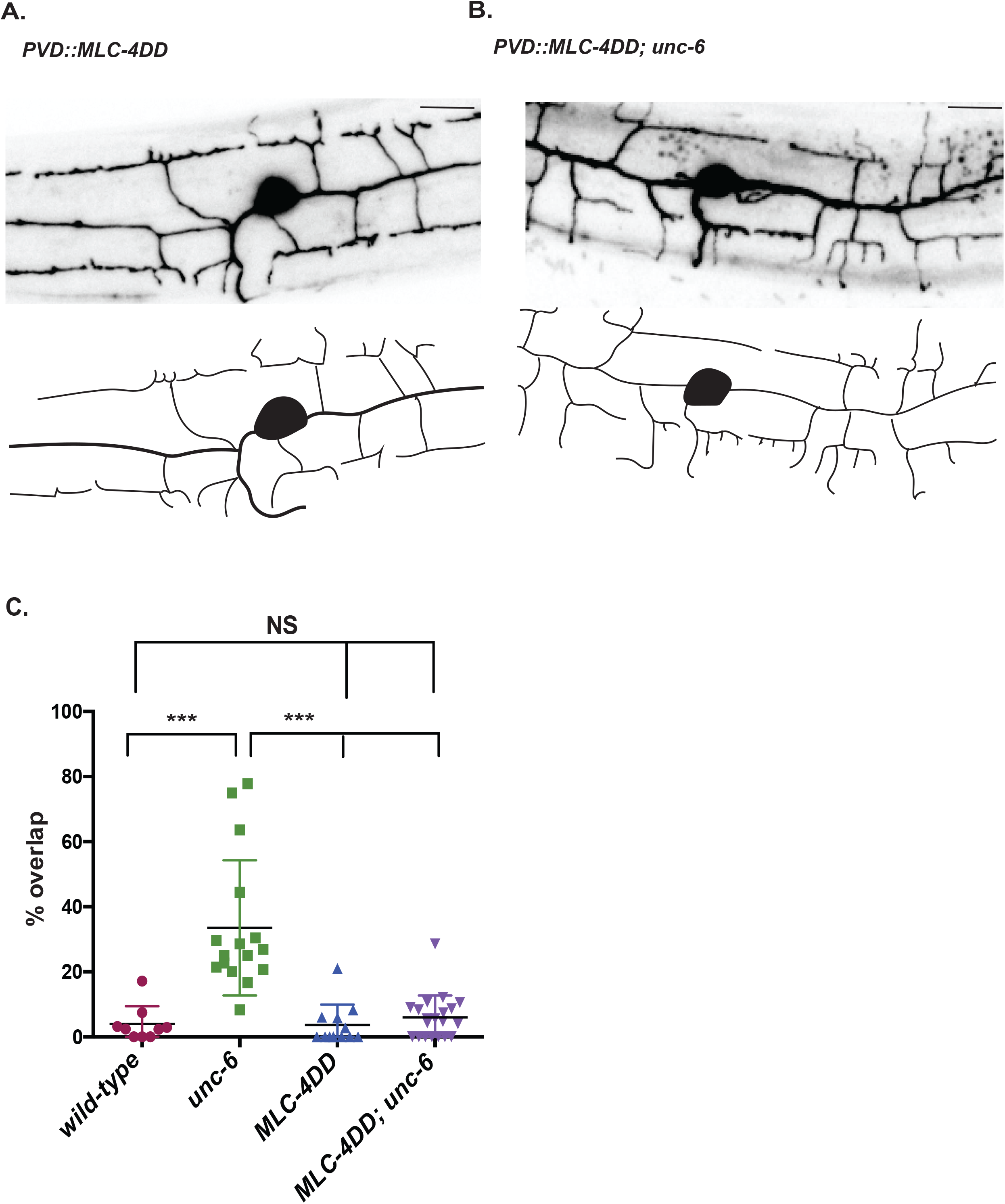
Phosphomimetic activation of myosin regulatory light chain, MLC-4, rescues the *unc-6* dendritic self-avoidance defect. (A-B) Representative images and tracings of PVD dendritic architecture in late L4 stage larvae expressing (A) PVD::MLC-4DD or (B) PVD::MLC-4DD; *unc-6(ev400).* (C) PVD expression of MLC-4DD rescues the *unc-6* mutant self-avoidance defect, ***p<0.0001, One way ANOVA with Tukey’s posthoc test. Mean and SEM are shown. n ≥ 9 animals per genotype. Quantified with PVD::mCherry. Scale bar is 10 μm. wild-type and *unc-6(ev400)* results from Figure 2D.

### Antagonistic pathways regulate dendrite outgrowth vs self-avoidance

Our time lapse imaging studies detected dynamic actin polymerization in growing PVD dendrites (Movie S3). This observation and the recent finding that PVD dendrites are truncated and misdirected by mutations that disable either a specific actin structural gene, *act-4*, or specific components (WRC and TIAM/GEF) that promote actin polymerization, suggest that dendritic outgrowth depends on actin assembly [26,27] (Figure S2D). Actin assembly is also likely required for dendrite retraction given results showing that genetic knockdown of a separate set of actin-binding proteins (i.e., UNC-34/Ena/VASP, MIG-10/Lpd, UNC-73/Trio, WSP-1/WASP, ARX-5/p21) (Fig. 3), blocks PVD self-avoidance but not dendrite outgrowth [37]. A shared role for actin polymerization in both outgrowth and retraction argues that the net length of PVD dendrites must depend on an intrinsic mechanism that regulates actin assembly to balance these competing effects.

Our results suggest that UNC-6/Netrin signaling promotes dendrite retraction in a mechanism involving F-actin assembly. PVD dendritic outgrowth is driven by a separate pathway in which DMA-1, a membrane protein expressed in PVD, interacts with a multicomponent receptor anchored in the adjacent epidermis [13,45–49]. Dendritic growth depends on binding of DMA-1 and the claudin-like protein, HPO-30, to specific regulators of actin assembly (i.e., WAVE regulatory complex, TIAM-1/GEF) in the PVD cytoplasm [26,27,50]. These interactions may be modulated by the Furin KPC-1 which antagonizes DMA-1 surface expression in PVD [45–47]. *kpc-1* activity is proposed to temporally weaken adhesion with the epidermis to facilitate changes in trajectory at specific stages in dendritic outgrowth [47,48]. For example, the tips of 3° branches typically execute a right-angle turn after the self-avoidance response to produce a 4° dendrite [11]. This morphological transition largely fails in *kpc-1* mutants in which 3° dendrites elongate abnormally and show severe self-avoidance defects due to over-expression of DMA-1 [47,48]. Thus, we considered the idea that self-avoidance is achieved by the combined effects of *kpc-1*-dependent downregulation of DMA-1 and a separate mechanism that antagonizes outgrowth and requires UNC-6/Netrin. We performed a series of genetic experiments to test this model.

The hypomorphic loss-of-function allele, *kpc-1(xr58)*, results in over-expression of DMA-1 and a consequent, robust self-avoidance defect with ~42% of sister 3° branches overlapping one another (Figure 9A, D) [46,47]. This defect is not observed, however, in *kpc-1/+* heterozygotes (Figure 9D). Similarly, self-avoidance in *unc-6/+* heterozygous animals is indistinguishable from wild type (Figure 9D). Thus, a 2-fold reduction in gene dosage for either *kpc-1* or *unc-6* does not perturb self-avoidance. We reasoned, however, that if expression of UNC-6/Netrin and KPC-1 were simultaneously reduced by half, then self-avoidance should be at least partially disabled since both *unc-6* and *kpc-1* normally function to antagonize dendrite outgrowth. To test this prediction, we constructed double heterozygous mutants of *kpc-1* and *unc-6* (e.g., *kpc-1/+; unc-6/+*) and quantified the self-avoidance phenotype. This experiment revealed that ~30% of adjacent sister 3° dendrites fail to self-avoid in *kpc-1/+; unc-6/+* animals which is significantly greater than that of either *kpc-1/+* or *unc-6/+* single mutants (Figure 9B, D). This genetic “enhancer” effect is consistent with the idea that both KPC-1 and UNC-6 function to limit growth of 3° dendrites. If this effect is due to elevated levels of DMA-1 arising from reduced KPC-1 activity in the *kpc-1/+; unc-6/+* background, then a concomitant diminution of DMA-1 activity should restore normal self-avoidance. We tested this idea by constructing the triple heterozygote, *kpc-1 +/+ dma-1; unc-6/+*, and detected robust suppression of the self-avoidance defect observed in *kpc-1/+; unc-6/+* animals (Figure 9C, D). Thus, these results suggest that DMA-1-mediated adhesion to the epidermis antagonizes UNC-6/Netrin-dependent dendrite retraction and that the relative strengths of these opposing pathways must be finely tuned to achieve the self-avoidance effect (Figure 9E). The key role of actin polymerization in both UNC-6/Netrin-mediated retraction and DMA-1-dependent outgrowth argues that this balancing mechanism likely regulates both modes of actin assembly.

**Figure 9.**
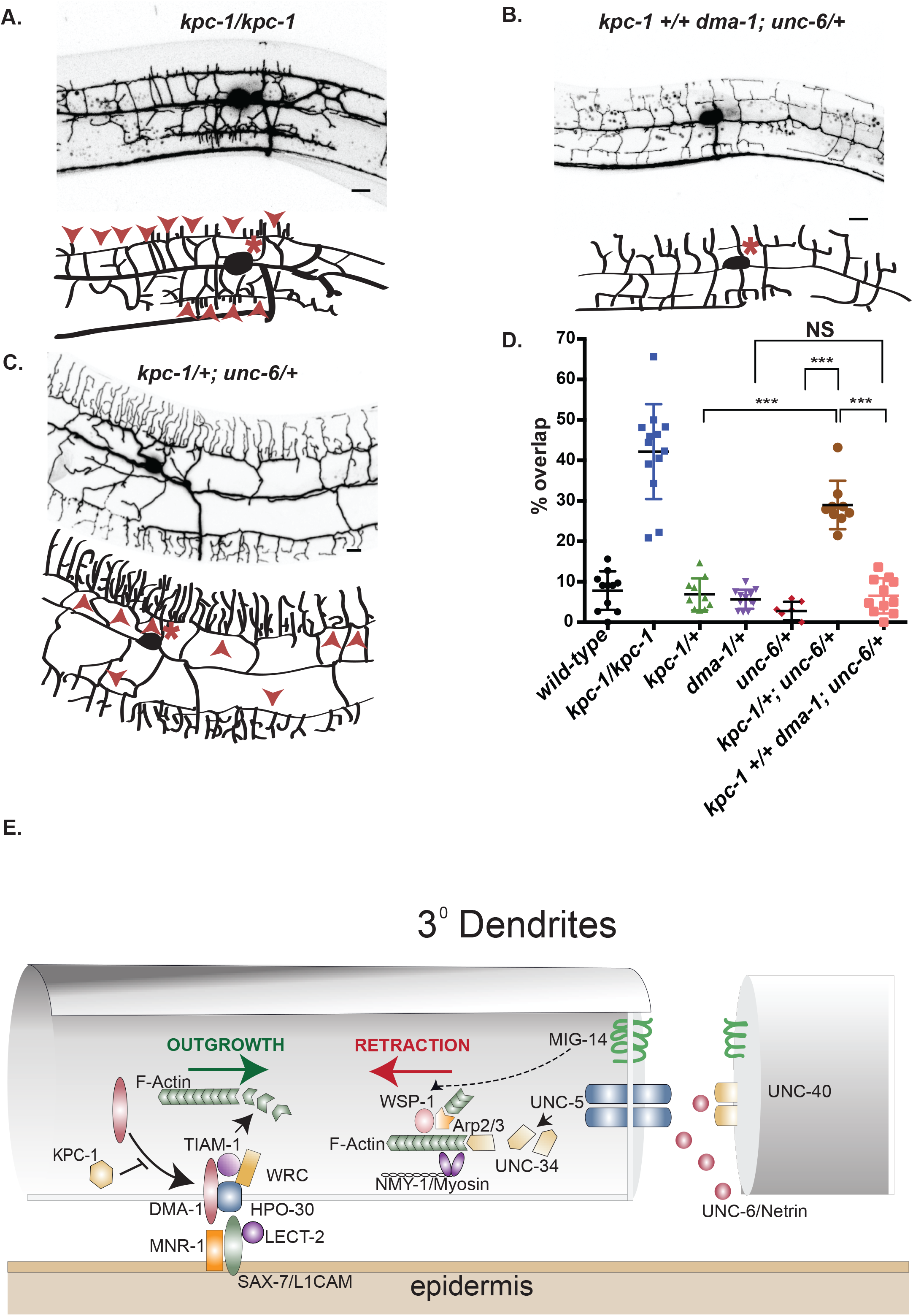
Opposing pathways regulate dendrite outgrowth versus retraction. Representative images and schematics of PVD in (A) *kpc-1/kpc-1*, (B) *kpc-1/+; unc-6/+* and (C) *kpc-1 +/+ dma-1; unc-6/+*. PVD soma (asterisks) and overlapping 3° dendrites (arrowheads) are noted. (D) Quantification of self-avoidance defects for each genotype. Note that all mutant alleles are recessive but that *kpc-1/+; unc-6/+* double heterozygotes show a strong self-avoidance defect that is suppressed in *kpc-1 +/+ dma-1; unc-6/+* triple heterozygotes. ***p<0.0001, one way ANOVA with Tukey’s posthoc test. Mean and SEM are indicated. *** p <0.0001, N >7 animals per genotype. PVD is labeled with *ser2prom3*::GFP for all images. Scale bar is 10 μm. Mutant alleles were: *kpc-1(xr58), unc-6(ev400)* and *dma-1(wy686).* (E) Model of opposing pathways that promote actin polymerization to define dendrite length. KPC-1 antagonizes surface expression of DMA-1 that interacts with an epidermal receptor composed of SAX-7/CAM, MNR-1 and LECT-2 and functions with HPO-30 to activate TIAM-1/GEF and the Wave Regulatory Complex (WRC) for actin polymerization and dendrite outgrowth. UNC-6/Netrin is captured by UNC-40/DCC for interaction with UNC-5 to activate UNC-34/Ena/VASP which functions with WSP-1/WASP and the Arp2/3 complex to drive F-actin assembly for NMY-1/myosin-dependent dendrite retraction in the self-avoidance response. (UNC-73/Trio and MIG-10/Lamellipodin are not shown to simplify the diagram, see Results). Homotypic interaction of MIG-14/Wntless in contacting 3° dendrites also drives retraction possibly through WSP-1.

## DISCUSSION

Dendrites arising from the same neuron typically do not overlap. This phenomenon of dendrite self-avoidance is universally observed and derives from an active mechanism in which contact between sister dendrites triggers mutual retraction [4,48]. As might be predicted for an event that depends on physical proximity, cell-surface associated proteins have been shown to mediate dendrite self-avoidance [37,49]. These include Dscam and protocadherins which are expressed in multiple alternative forms to distinguish self from non-self among contacting dendrites [5–7]. Self-avoidance for certain neurons can also depend on the interaction of diffusible cues with their cognate receptors [8,49]. For example, we have shown that the axon guidance signal, UNC-6/Netrin, and its receptors, UNC-5 and UNC-40/DCC, function together to mediate self-avoidance for PVD sensory neurons in *C. elegans*. We have proposed that UNC-40/DCC captures UNC-6/Netrin on the membrane surface to facilitate repulsion through physical interaction with UNC-5 on PVD dendrites [9] (Figure S1D). Little is known, however of the downstream effectors of dendrite retraction in the self-avoidance response. Here we report new results that are consistent with a model in which UNC-6/Netrin promotes actin-polymerization to propel dendrite retraction. Our finding that non-muscle myosin is also involved resolves the paradoxical idea that UNC-6/Netrin-dependent actin polymerization shortens rather than lengthens PVD 3° dendrites. We propose that non-muscle myosin engages the F-actin cytoskeleton at the tips of PVD dendrites to drive retraction.

### Regulators of actin polymerization are required for dendrite self-avoidance

The proposed role for actin polymerization in the self-avoidance response derives from our finding that multiple regulators of F-actin assembly are necessary for efficient dendrite retraction. For example, a mutation that disables UNC-34, the *C. elegans* homolog of Ena/VASP, disrupts PVD self-avoidance. Ena/VASP promotes actin polymerization by preventing capping proteins from blocking the addition of actin monomers to the plus-end of elongating actin filaments. In axon growth cones, Ena/VASP localizes to the tips of actin bundles in growing filopodia [12,36]. We observed punctate Ena/VASP localization at the tips of PVD dendrites and active trafficking from the cell soma (Figures 4 and S3). Our results showing that an *unc-34* mutant suppresses the retarded dendrite outgrowth phenotype arising from the constitutively active MYR::UNC-5 (Figure 3) argues that *unc-34* functions downstream to mediate UNC-6/Netrin-dependent self-avoidance. Additional genetic evidence indicates that WSP-1/WASP also acts in a common pathway with UNC-6/Netrin (Figure 3). The established role for WASP of activating the Arp2/3 complex [50] is consistent with our finding that RNAi knockdown of *arx-5/p21*, a conserved Arp2/3 component, partially disables the self-avoidance mechanism (Figure 3). Dual roles for UNC-34 and WSP-1 in actin polymerization, as suggested here, are consistent with previous work indicating that Ena/VASP can activate Arp2/3 through direct interaction with WASP proteins [31,51–53]. Thus, taken together, our results suggest that Ena/VASP and WSP-1 coordinate the creation of branched actin networks at the tips of retracting PVD dendrites in the UNC-6/Netrin dependent self-avoidance response (Figure 9E).

A role for Ena/VASP in dendrite retraction, as have we have proposed here, parallels earlier findings of a necessary function for Ena/VASP in growth cone repulsion. For example, axon guidance defects arising from ectopic expression of UNC-5 in *C. elegans* touch neurons requires UNC-34/Ena/VASP [54], a finding confirmed by our observation that the dendritic shortening phenotype induced by MYR::UNC-5 is prevented by a loss-of-function mutation in *unc-34*. Ena/VASP is also required for axon repulsion mediated by Slit and its receptor Robo [20,55,56]. Although Ena/VASP function can be readily integrated into models of axonal attraction, the well-established role of Ena/VASP in F-actin assembly has been more difficult to rationalize in mechanisms of axonal repulsion which are presumed to involve actin depolymerization [12,57]. A partial explanation for this paradox is provided by a recent study showing that Slit, acting through Robo, induces transient filopodial outgrowth in migrating axonal growth cones and that this effect requires Ena/VASP [56]. The functional necessity for filopodial growth in this mechanism is unclear, but the central role of Ena/VASP-induced actin assembly in the overall repulsive response to Slit mirrors our results suggesting that dendrite retraction in PVD neurons requires UNC-34/Ena/VASP and F-actin assembly.

Our studies also detected necessary roles in self-avoidance for MIG-10/LPD which has been previously shown to function with UNC-34/Ena/VASP to promote F-actin polymerization and UNC-73/Trio, a Rho/Rac GEF that acts in UNC-6/Netrin pathways to regulate the actin cytoskeleton [15,19,20,27]. Altogether, our genetic analysis identified five known effectors of actin polymerization (UNC-34/Ena/VASP, WSP-1/WASP, Arp2/3 complex, MIG-10/Lpd, UNC-73/Trio) that mediate the self-avoidance response but which are not required for 3° branch extension during outgrowth (Figure 3, S2).

### Non-muscle myosin II mediates dendrite retraction in the self-avoidance response

Our genetic analysis detected a cell-autonomous role for NMY-1/non-muscle myosin II in dendrite self-avoidance (Figure 6). Because our results revealed F-actin at the tips of PVD dendrites (Figure 5), we suggest that NMY-1 mediates retraction by interacting with F-actin to drive a contractile mechanism. For example, NMY-1 could accelerate retrograde flow, a non-muscle myosin-dependent effect that also drives filopodial retraction in growth cones and at the leading edge of migrating cells [58,59]. Alternatively, NMY-1 could shorten PVD dendrites by inducing the reorganization of the nascent actin cytoskeleton into a more compact structure [60]. Phosphorylation of myosin regulatory light chain (RLC) activates non-muscle myosin [43] and we showed that MLC-4DD, a phosphomimetic, and, thus, constitutively active form of the *C. elegans* homolog of RLC, results in delayed outgrowth of PVD 3° dendrites (Figure 7). The retarded dendrite outgrowth resulting from chronic activation of MLC-4 suggests a model in which NMY-1 function is triggered by contact between sister dendrites and subsequent UNC-6/Netrin signaling. A downstream function for non-muscle myosin II in the UNC-6/Netrin pathway is consistent with our finding that the Unc-6 self-avoidance defect is rescued by MLC-4DD (Figure 8). Parallel roles for these components in axon guidance are suggested by recent results showing that non-muscle myosin II is required for Slit and Netrin-dependent midline avoidance in the developing vertebrate spinal cord [61]. Similarly, non-muscle myosin II mediates Semaphorin-induced axonal repulsion. The mechanism of Semaphorin action in this case also involves the assembly of actin bundles in the axon shaft presumably to facilitate non-muscle myosin-driven retraction [62–64]. Thus, our findings suggest that key drivers of axonal repulsion including nascent actin assembly and non-muscle myosin may also mediate dendrite retraction in the self-avoidance response.

RLC phosphorylation activates non-muscle myosin II by releasing myosin monomers for assembly into bipolar structures or myosin “stacks” that interact with actin filaments to induce translocation [43,65]. Although our results are consistent with a necessary role for non-muscle myosin in dendrite retraction, the tips of PVD 3° dendrites are likely too narrow (~50 nm) [10] to accommodate myosin stacks (~300 nm) [65,66]. We suggest that the actin-branching function of Arp2/3, which we have shown mediates PVD dendrite retraction, could induce transient expansion of the dendritic tip to allow assembly of myosin stacks. In this model, actin-polymerization involving Arp2/3 could effectively limit ectopic NMY-1 activation in the dendritic shaft by restricting the assembly of myosin stacks to the tips of contacting sister dendrites. Alternatively, recent evidence indicates that myosin can also induce contraction in its monomeric or unpolymerized form [67] which should have ready access to the dendrite tip. This idea is consistent would our observation that GFP-tagged NMY-1 is active (e.g., rescues the self-avoidance defect of *nmy-1*) and did not display the punctate appearance characteristic of myosin stacks [65] but is diffusely localized throughout the PVD cytoplasm (Figure 6B).

### Parallel-acting pathways regulate dendrite self-avoidance

Although self-avoidance is perturbed in UNC-6/Netrin pathway mutants, a significant fraction (~70%) (Figure 3D) of PVD 3° dendrites show apparently normal self-avoidance behavior [9]. One explanation for this observation is provided by the recent discovery of an independent pathway, mediated by MIG/14/Wntless, that is necessary for self-avoidance in an additional fraction of PVD 3° dendrites [37]. Parallel acting pathways also mediate self-avoidance in mammalian Purkinje neurons where slit-robo signaling acts in concert with protocadherins [6,8]. The likelihood of additional effectors of self-avoidance is also suggested by the incompletely penetrant effects of Dscam mutants in Drosophila sensory neuron self-avoidance [68]. Thus, our results in *C. elegans* mirror findings in other organisms which together suggest that multiple mechanisms have evolved to insure dendrite self-avoidance.

### The actin cytoskeleton is highly dynamic in growing and retracting PVD dendrites

We used time-lapse imaging with LifeAct::GFP, a live-cell marker for actin {Riedl:2008gw}, to monitor actin PVD dendrites. The LifeAct::GFP signal is typically brighter near the distal ends of growing branches in comparison to interstitial regions (Figure 5) and is highly dynamic (Movie S3). The distal localization and rapidly fluctuating LifeAct::GFP signal in PVD dendrites resembles recently reported “actin blobs” that actively migrate in highly branched Drosophila sensory neurons where they appear to presage points of branch initiation [69]. A strong LifeAct::GFP signal at the tips of emerging PVD dendrites is also consistent with genetic evidence that actin polymerization is required for dendrite outgrowth [24,25]. Although our genetic results (Figure 3) suggest a model of dendrite retraction in which nascent actin polymerization is necessary for the self-avoidance response, our measurements did not detect a quantitative increase in LifeAct::GFP in retracting vs growing 3° dendrites as previously reported by others [37]. This result suggests that the overall level of actin polymerization is high during both outgrowth and retraction but may be incorporated into distinct structures, regulated by distinct groups of effectors (see below) that our imaging methods do not resolve.

### Antagonistic pathways regulate dendrite outgrowth vs retraction

Dendritic architecture is defined by the combined effects of outgrowth which expands the arbor vs retraction which limits the size of the receptive field. This interaction of positive and negative effects is readily observed in the development of 3° PVD dendrites. Initially, 3° dendrites emanating from adjacent menorahs grow out along the body axis until contacting one another and then retract to avoid overlap [9,11]. Outgrowth in this case is promoted by a multicomponent receptor-ligand complex that mediates adhesive interactions with the adjacent epidermis [70–73] (Figure 9E). When this pathway is dysregulated by over-expression of the DMA-1 receptor in PVD neurons, for example, 3° dendrites continue to adhere to the epidermis and overgrow one another [47,74]. 3° branch self-avoidance also fails in mutants that disable UNC-6/Netrin signaling which we have shown promotes dendrite retraction [9]. Thus, these results suggest that the net length and placement of each 3° branch depends on the balanced effects of locally acting signals that either extend (e.g., DMA-1) or shorten (e.g., UNC-6/Netrin) the dendrite. We confirmed this idea in genetic experiments that detected strong dose-sensitive interactions between these opposing pathways. For example, a 50% reduction of UNC-6/Netrin expression in a heterozygous *unc-6/+* mutant does not perturb self-avoidance. Similarly, a genetic mutant (*kpc-1/+)* that partially elevates DMA-1 also shows normal 3° branch outgrowth. The combination of both mutations in the double heterozygote, *unc-6/+; kpc-1/+*, however, produces a strong self-avoidance defect (Figure 9). Our genetic results identified multiple regulators of actin polymerization (UNC-34/Ena/VASP, WSP-1/WASP, ARX-5/p21, MIG-10/Lamelipodin, UNC-73/Trio) that are required for self-avoidance (Figure 3). We thus propose that UNC-6/Netrin promotes actin polymerization to effect dendrite retraction and that non-muscle myosin II interacts with a nascent actin cytoskeleton to translocate each 3° dendrite away from its neighbor. F-actin is also abundant and highly dynamic in growing PVD dendrites. Notably, a distinct set of F-actin promoting factors (WRC, TIAM/GEF) interacts with the DMA-1 receptor complex to drive PVD dendritic growth (Figure 9E) [26,27]. Additional genetic evidence suggests that the WRC could also function in self-avoidance [37]. Thus, the challenge for future studies is to elucidate the cell biological machinery through which opposing pathways manipulate the actin cytoskeleton to effect either dendrite growth or retraction.

## MATERIALS AND METHODS

### Strains and genetics

All the strains used were maintained at 20°C and cultured as previously described (Brenner, 1974). We used the N2 Bristol strain as wild-type.

Additional strains used in this study: NC1404 [*wdIs52* (*pF49H12.4*::GFP)], NW434 [*unc-6(ev400)X*;*wdIs52(pF49H12.4::GFP)*], TV17,200 [*kpc-1(xr58); wyIs592(pser2prom3::GFP)*] [47], TV9656 [*dma-1(wy686); unc-119(ed3)III; wyEx3355(pser2prom3::GFP+odr-1::RFP)*], CX1248 [*KyEx3482*(*pdes-2::TBA-1::mCherry+coel::RFP*)] {Maniar:2011gx}, CLP928 [*twnEx382(Pser2.3::LifeAct::EGFP, Pser2.3::mCherry, Pgcy-8::gfp)*] [37].

Additional strains generated for this study: NC2580 *unc-34(gm104)V*; *wdIs52*, NG324 [*wsp-1(gm324)*; *wdIs52*], *+/szT1 lon-2(e678) I;* HR1184 [*nmy-1(sb115) dpy-8(e130)/szT1 X*], NC2726 [*nmy-1(sb113)*;*wdIs52*], *unc-73(rh40)*]; *wdIs52*.

NC3284 [*wdEx1011* (*pF49H12.4*::myrUNC-5::GFP)], was produced by microinjection of plasmids pLSR18 (*pF49H12.4*::myrUNC-5::GFP), *pmyo-2*::mCherry and pCJS04 (*pF49H12.4*::mCherry).

NC3048 [*wdIs98* (*pF49H12.4*::ACT-1::GFP)] was produced by microinjection of plasmids pLSR03 (*pF49H12.4*::ACT-1::GFP) and *pceh-22*::GFP and integrated as previously described (Miller and Niemeyer, 1995).

NC3085 [*wdEx967* (*pF49H12.4*::mCherry::UNC-34)] was obtained by microinjection of plasmids pCJS78(*pF49H12.4*::mCherry::UNC-34) and *pmyo-2*::mCherry.

NC3090 [*wdEx972* (*pF49H12.4*::LifeAct::GFP)] was acquired by microinjecting plasmids pLSR06 (*pF49H12.4*::LifeAct::GFP) and *pmyo-2*::mCherry.

NC3029 [(*pF49H12.4*::*nmy-1(+)* + *pF49H12.4*::*nmy-1(-)*)] was obtained by microinjection of pLSR09 (*pF49H12.4*::*nmy-1(+)*), pLSR10 (*pF49H12.4*::*nmy-1(-)*), pmyo-2::mCherry and pCJS04 (*pF49H12.4*::mCherry)

NC3087 [*wdEx969* (*pF49H12.4*::GFP::Linker::NMY-1)] was produced by microinjecting plasmids pLSR11 (*pF49H12.4*::GFP::Linker::NMY-1), pCJS04 (*pF49H12.4*::mCherry) and pmyo-2::mCherry.

*wdEx1009* (*pF49H12.4*::GFP::Linker::MLC-4DD) [NC3283] was obtained by microinjection of plasmids pLSR17 (*pF49H12.4*::GFP::Linker::MLC-4), pCJS04 (*pF49H12.4*::mCherry) and *pmyo-2*::mCherry.

### Molecular Biology

#### pLSR11 (*pF49H12.4::*GFP::Linker::NMY-1::*unc-10* 3’UTR)

We used the Clonetech infusion HD cloning kit (Cat # 638910) to build this plasmid. The *nmy-1* genomic region was amplified from pACP01 (*pF49H12.4::*NMY-1::mCherry) with overlapping regions corresponding to pCJS95 (*pF49H12.4::*GFP::CED-10::*unc-10* 3’UTR). We also PCR amplified pCJS95 with primers overlapping the *nmy-1* genomic sequences to swap CED-10 with NMY-1. We then used the NEB site-directed mutagenesis kit to insert a 24-nucleotide glycine rich linker (see Table 1) [65]. The resultant pLSR11 plasmid was sequenced to confirm that the linker was inserted between the GFP sequence and *nmy-1*. See table 1 for primer sequences.

**Table 1.**
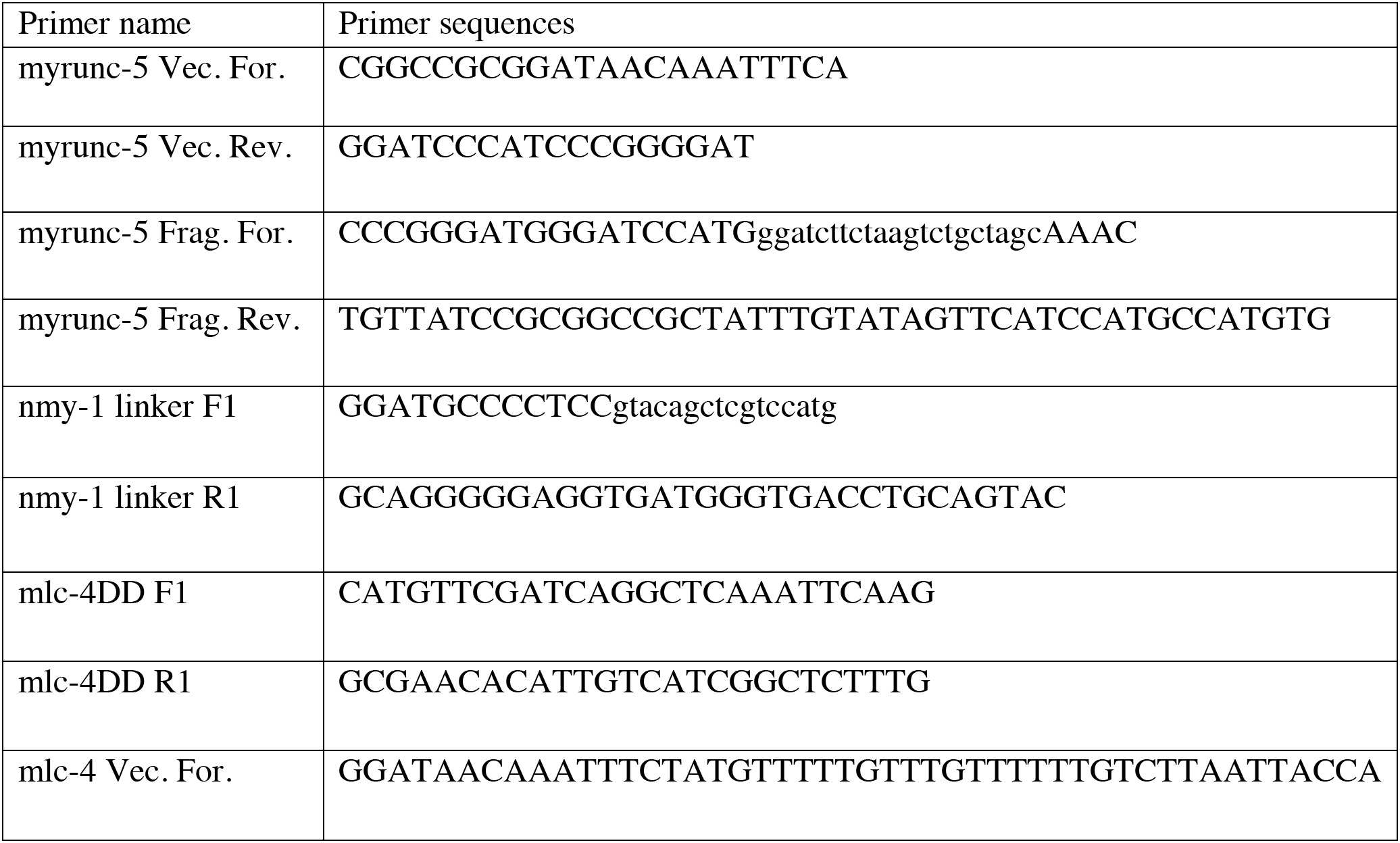
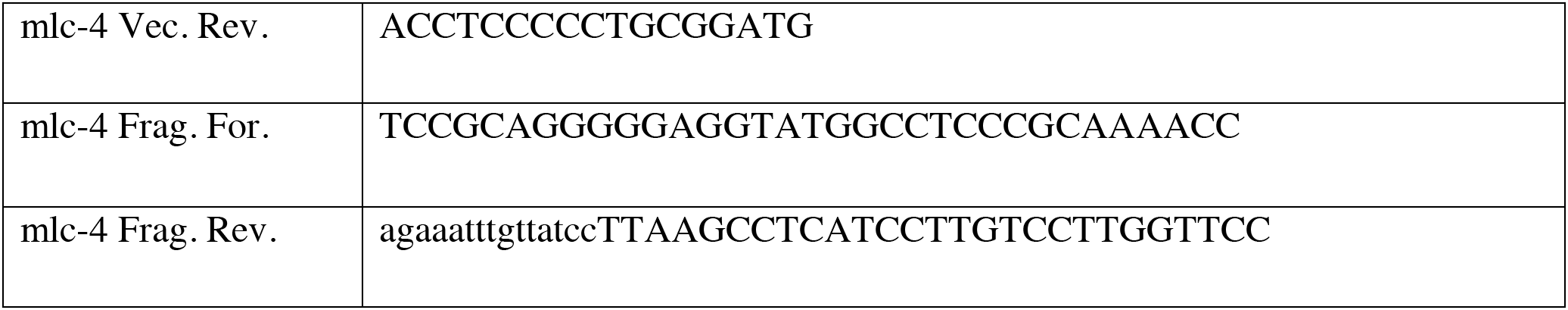

#### Constructing pLSR17 (*pF49H12.4*::GFP::Linker::MLC-4DD)

We used the Clonetech infusion HD cloning kit (Cat # 638910) to construct this plasmid. The *mlc-4* genomic region was amplified from wild-type (N2) DNA using primers with overlapping adapters complementary to pLSR11 (*pF49H12.4*::GFP::Linker::NMY-1). pLSR11 was amplified with primers containing adapter sequences complementary to *mlc-4*. The amplified fragments were combined for the infusion cloning to replace *nmy-1* with *mlc-4*. We then used the NEB site directed mutagenesis kit to change MLC-4 codons for Serine 18 and Threonine 17 to nucleotide sequences that encode aspartatic acid (D). The resultant pLSR17 plasmid was confirmed by sequencing.

#### Confocal microscopy and image analysis

Animals were immobilized with 15 mM levamisole/ 0.05% tricaine and mounted on 2% agarose pads in M9 buffer as previously described [11]. Z stack images were collected using a Nikon TiE inverted fluorescent microscope equipped with an A1R point scanning confocal and a Plan Apo 40X, 1.3NA oil immersion objective or on a Leica TCS SP5 confocal microscope with a 40X or 63X objective. Z stacks were collected at either 0.5 or 1μm depth from the position of ventral nerve cord to the top of PVD cell body. Maximum projection images were collated with either LAS-AF (Leica) or with NIS elements software (Nikon). Dendrite length was measured using the line tool in Fiji/image J. Co-localization images were assembled using image J. Z stack images that were collected using the LAS-AF or NIS elements software was imported into ImageJ/Fiji for further analysis.

#### iSIM super-resolution imaging and analysis

Animals were immobilized with 0.4 mM levamisole and mounted on 10% agarose pads in M9 buffer. Z stack images were captured using a Hamamatsu sCMOS camera on a iSIM microscope (VisiTech International) equipped with a 100X, 1.45 NA objective. Images were deconvolved and processed using Metamorph and Fiji/ImageJ.

#### Imaris software image analysis

Time-lapse movies from the iSIM microscope were deconvolved with metamorph. The spot tracking algorithm (Imaris) was used to track UNC-34::mCherry particles of minimum diameter of 0.3 µm. Displacement refers to the distance (μm) between the location of an UNC-34::mCherry particle at a given time versus its starting position at t = 0.

#### Spinning disk confocal microscopy

Animals were immobilized with 0.4mM levamisole and mounted on 10% agarose pads in M9 buffer. Volumes over time were captured using an automated TiE inverted fluorescence microscope platform with an X1 spinning disk head (Yokogawa) and DU-897 EM-CCD (Andor Technology), piezo-electric Z stage (Mad City Labs) and Apo TIRF 100x 1.49 NA oil immersion objective (Nikon Instruments, Inc.). Analysis of fluorescence intensity was accomplished using NIS-Elements and Fiji/Image J software.

#### Quantification of fluorescence intensity in growing vs. non-growing dendrites

L2-L3 larvae were immobilized with 0.4mM levamisole and mounted on 10% agarose pads in M9 buffer for imaging with the spinning disk confocal microscope. Z-stacks of PVD dendrites were imaged at either 1 min or 1.30 minute intervals with the 100X 1.49 NA oil immersion objective. PVD dendrites (2º, 3º and 4º) undergoing outgrowth with a succession of rapid extensions and retractions during the length of the time lapse videos were analyzed. NIS elements was used for measurements of LifeAct::GFP from a 1µm ROI at the tips of growing dendrites. In the same videos, fluorescence intensity was measured for an identical 1µm ROI of PVD branches (1º, 2º and 3º) that were not undergoing outgrowth. Fluorescence intensity measurements for the outgrowth and no-outgrowth datasets were normalized against the maximum overall LifeAct::GFP value (Figure 5).

#### Statistics

Student’s t test was used for comparisons between 2 groups and ANOVA for comparisons between three or more groups with either Tukey posthoc or Bonferroni correction for multiple comparisons. Kolmogorov-Smirnov test was performed for comparisons between frequency distributions of 3° dendrite length and gaps between 3° dendrites (Figures 2C and 8F, G).

## ACKNOWLEDGMENTS

We thank C. Bargmann, K. Shen, J. Plastino, Dr. T.J. Kubiseski, E. Lundquist, P. Mains and H. Buelow for strains and reagents. We thank Chun-Liang Pan and Chien-Po Liao for sharing their work, reagents and advice prior to publication, Dylan Burnette and members of his laboratory, Aidan Fenix and Nilay Taneja, for helpful suggestions and members of the Miller lab for comments on the manuscript. Spinning disk images were obtained in the Nikon Center of Excellence at Vanderbilt University. We thank Fernando Delaville and Ralph Bauer from BioVision Technologies for their help in acquiring time-lapse movies and images on the iSIM microscope. This work was supported by NIH Grants R01 NS079611 to D. M. M, F31 NS071801 to C.J.S and F31 NS49743 to J. D. W. Some of the strains used in this work were provided by the *C. elegans* Genetics Center, which is supported by the US National Institutes of Health (NIH) National Center for Research Resources. L.S., C.J.S and J.D.W. performed experiments with advice from D.M.M., M.J.T. provided imaging expertise and advice; B.A.M helped with spinning disk imaging and edited videos, L.S. and D.M.M. wrote the paper with input from coauthors.

## SUPPORTING INFORMATION

### Airy Scan imaging and analysis

Animals were immobilized with 15mM levamisole/0.05% tricaine and mounted on 10% agarose pads in M9 buffer. Z stacks were captured using a Plan Apo 63x, 1.40 NA objective on a Zeiss LSM880 inverted fluorescence microscope outfitted with Airy Scan technology (Carl Zeiss Microscopy). Images of PVD::LifeAct::GFP and PVD::mCherry::UNC-34 (Figure S3 A-B) were processed to achieve Airy Scan (1.7x) super resolution and cropped using Fiji/ImageJ.

### TIRF microscopy and analysis

Animals were immobilized with 15mM levamisole/ 0.05% tricane and mounted on 2% agarose pads in M9 buffer. The coverslip placed on top of the agarose pad encasing the immobilized worms is 1mm in thickness. Time-lapse datasets were acquired using a Nikon TiE inverted fluorescence microscope outfitted with a TIRF illuminator, Apo TIRF 100x, 1.49NA oil immersion objective, and NIS-Elements software (Nikon Instruments, Inc.). The images for time lapse videos of PVD::mCherry::UNC-34 (Figure S3C) were captured in a single plane at 15 frames/sec and the TIRF angle was adjusted for oblique illumination of 3° PVD dendrites. Datasets were imported into FIJI/ImageJ for further analysis.

### Imaging and quantification of self-avoiding 3° dendrites

#### Condition 1

(See Figure S4A-C) L3 animals of strain CLP928 *twnEx382[Pser2.3::LifeAct::EGFP, Pser2.3::mCherry, Pgcy-8::gfp]* [37] were immobilized for time-lapse imaging with 1 µl of agarose beads (Polybead carboxylate microspheres, Polysciences, Catalog # 15913-10, 0.05 µm) and mounted on a 10% agarose pad. Z-stacks of PVD dendrites (0.5 um depth, 5 slices) were imaged at both 35 second intervals with a 40X objective in a Nikon spinning disk confocal microscope. Contact between the 3° dendrites was established as a function of max feret (maximum distance between 2 points). Max feret (in pixels) was measured in the mCherry channel (cystosolic marker) with NIS elements and contact defined as the time-point at which the max-feret approaches zero. LifeAct::GFP and mCherry signals were measured for a ROI of 2 µm at the tips of the left and right dendrites in a single Z plane. Left and right fluorescence intensities were summed for comparison to a control ROI (2 µm) in an adjacent non-contacting region. The change in fluorescence intensity (∆F) for each contact and control region was calculated as the difference between the signal at contact (F_t_) and the fluorescence value at two time points prior to contact (F_t-2_) normalized to F_t-2_ or ∆F = (F_t_ - F_t-2_/F_t-2_ [37]. These values were computed for contact and control ROIs for LifeAct::GFP and mCherry. Statistical tests used for comparison between the datasets (LifeAct::GFP contact vs control, mCherry contact vs control, LifeAct::GFP contact vs. mCherry contact) were 2-way ANOVA with Bonferroni correction and multiple t-tests with a False Discovery Rate of 1%. Both these tests yielded similar results. No significant differences were detected in the LifeAct::GFP signal in contacting vs non-contacting regions of 3° dendrites.

#### Condition 2

Late L3 larvae of P*F49H12.4::LifeAct::GFP; PF49H12.4::mCherry* were immobilized as above for imaging in the spinning disk confocal microscope. Z-stacks of PVD dendrites were imaged at either 1 or 1.30 minute intervals with a 100X objective (See Figure S4D). Contact is defined by the point at which the max feret (maximum distance between 2 points) between adjacent PVD 3º dendrites in the mCherry channel approached zero. For comparisons of LifeAct::GFP signals at the tips of 3º dendrites before and during contact, fluorescence intensities were determined from a 5 µm ROI centered at the point of contact and from a contiguous control (non-contact) 5 µm ROI positioned either to the left or right of the contact region (Figure 6F). The change in fluorescence intensity (∆F) for each contact and control region was calculated as the difference between the signal at contact (F_t_) and the fluorescence value at two time points prior to contact (F_t-2_) normalized to F_t-2_ or ∆F = (F_t_ - F_t-2_)/F_t-2_ [37]. For all measurements, the cytoplasmic mCherry marker was monitored to measure the Max Feret (in this case, maximum distance between two points) between the tips of adjacent 3º dendrites and the point of contact determined when the Max Feret approaches zero. Statistical comparisons (N = 8) were performed with a 2-way ANOVA with Bonferroni correction (Figure S4D). For representative traces shown in Figure S4, Nikon NIS Elements was used to obtain LifeAct::GFP and mCherry fluorescence measurements from a 1 µm ROI at the tips of the left and right 3º dendrites undergoing contact. Fluorescence intensity measurements of LifeAct::GFP and mCherry for the opposing dendrites were normalized against the maximum value for each fluorophore and plotted vs time.

**Supplemental Figure 1: UNC-6/Netrin signaling mediates sister dendrite self-avoidance.**

(A-B) Representative images and tracings of PVD sensory neurons in wild-type (A) and *unc-6* (B) L4-stage larvae. PVD morphology was visualized with PVD::GFP or PVD::mCherry markers. PVD cell body marked with an asterisk. (A) 1°, 2°, 3° and 4° dendritic branches and single axon (blue arrowhead) are denoted. (B) Red arrowheads point to overlaps between adjacent 3° dendrites that failed to self-avoid in *unc-6* mutants. Scale bars are 5 μm. C) Summary of self-avoidance in 3° PVD dendrites denoting growth, contact and retraction. (D) Model of self-avoidance mechanism involving reciprocal contact-dependent repulsion mediated by UNC-6/Netrin and its receptors, UNC-40/DCC and UNC-5 [9].

**Supplemental Figure 2. Mutations in *unc-34/Ena/VASP* and *unc-73/Trio* disrupt PVD dendrite morphology.**

Quantification of 1° dendrite outgrowth and (B) number of 2° dendrites in wild-type, *unc-34*, *unc-73*, *wsp-1* and PVD::mCherry::UNC-34*;unc-34*. Defective 1° dendrites show altered alignment and/or extension defects. Mutations in *unc-34* and *unc-73* but not *wsp-1* result in defective 1° dendrite outgrowth and fewer 2° dendrites. Note that expression of UNC-34::mCherry in PVD restores 3° self-avoidance (Figure 3C) but does not rescue 1° and 2° dendritic defects which suggests that UNC-34 function could be required in other cell types for normal PVD 1° and 2° outgrowth, *p=0.02, **p=0.003, ***p<0.001, Fisher’s exact and 2-way ANOVA with Tukey’s correction for multiple comparison. (C) Representative images of PVD morphological defects in an *unc-34* mutant. Note premature termination of 1° dendrites (red arrowheads). (D) Representative image and drawing of PVD branching defects in a *tiam-1/GEF* mutant at the L4 stage. Scale bars are 10 μm. Mutant alleles are *unc-34(gm104), unc-73(rh40)*, *wsp-1 (gm324)* and *tiam-1(tm1556)*.

**Supplemental figure 3. Localization and trafficking of PVD:: mCherry ::UNC-34 puncta in PVD using Airy Scan and TIRF microscopy.**

(A) Zeiss Airyscan images of PVD simultaneously labeled with PVD::LifeAct::GFP (green) and PVD::mCherry::UNC-34 (Magenta). Merge shown on right. Insets denote PVD dendrites (B). White arrows indicate 2°, 3°, and 4° dendrites. Scale bar is 10 μm. (C) Kymographs of PVD::mCherry::UNC-34 in PVD 1° dendrite captured in the Nikon TIRF microscope. Anterograde (white) and retrograde (red) movements of mCherry::UNC-34 puncta are denoted.

**Supplemental figure 4. Actin dynamics during dendrite self-avoidance**

Schematic (left) of adjacent 3° dendrites and representative time lapse series (35 sec intervals) of PVD::mCherry and PVD::LifeAct::GFP in 3° dendrites showing self-avoidance response (Condition 1). Point of contact is indicated by an asterisk and period of contact denoted with dashed blue outline. Scale bar is 10 μm. (B) Schematic depicting ROIs for fluorescence intensity measurements at the tips of adjacent (left vs right) 3° dendrites that undergo contact and at an adjacent non-contact (control) region. (C) Graphical representation of the change in fluorescence intensity at each time point vs that of 2 time intervals (t-2) before contact (t = 0); Measurements were normalized to fluorescence intensity at t-2 or (F_t-2_) [37]. Normalized fluorescence intensity values were plotted against time (min) for LifeAct::GFP contacting 3° dendrites vs non-contact, and were compared by 2-way Anova with Bonferonni correction, p = 0.54 (N=14). (D) Time lapse series collected with 100X objective at 1-1.3 minute intervals (N = 8) (Condition 2). The values for LifeAct::GFP at contact vs non-contacting region of 3° dendrites at each time-point were compared by 2-way Anova with Bonferonni correction, p = 0.74, NS (Not Significant).

**Supplemental Figure 5: Actin dynamics during dendrite self-avoidance**

(A-D) Fluorescent intensity traces wild-type PVD dendrites during 3° dendrite self-avoidance. Fluorescent intensity measurements of PVD::LifeAct::GFP and PVD::mCherry were acquired at either 1 min (A-B) or 1.3 min (C-D) intervals from a 1µm region at the tips of the growing and contacting left and right 3° dendrites (Late L3-L4 animals). Measurements were normalized against the maximum intensity during a ~10 minute interval that includes at least one contact event. The gap between the left and right 3° dendrites undergoing contact was determined from the cytoplasmic mCherry signal and plotted against time. The period of contact (vertical shading) corresponds to a minimum value for the gap (0-0.14µm) between contacting left and right 3° dendrites. (E) Representation of 3° dendrites with color schemes to indicate left and right 3° dendrites and 1µm regions quantified.

**Movie S1: Time-lapse video representing the dynamic movement of 3° dendrites in a wild-type background.** Images of PVD::GFP were recorded (late L3-early L4) at 15 sec intervals with a 100X objective in a Nikon spinning disk microscope (see Materials and Methods). The length of the video is 240 seconds. The movie was edited using NIS elements software with Bayes advanced denoising. Green arrowheads denote the tips of 3° dendrites. Scale bar is 25 µm.

**Movie S2: Time-lapse video of 3° dendrites in a PVD::MYR::UNC-5.** Images of PVD::GFP in the PVD::MYR::UNC-5 strain were recorded (late L3-early L4) at 15 sec intervals with a 100X objective in a Nikon spinning disk microscope (see Materials and Methods). The entire length of the video is 12 minutes. Boxes denote tips of 3° dendrites showing limited outgrowth. The movie was edited using NIS elements software with Bayes advanced denoising. Scale bar is 10µm.

**Movie S3: Time-lapse video showing the dynamicity of LifeAct::GFP during in 3° PVD dendrites**. Images of PVD::mCherry (left) and PVD::LifeAct::GFP (right) and were collected at 1.3 min intervals with a 100 X objective in a Nikon spinning disk microscope (see Materials and Methods). The entire length of the video is 60 minutes. The movie was edited using NIS elements with automatic deconvolution. Scale bar is 5 µm. Spherical objects are autofluorescent granules.

**Movie S4: Time-lapse video showing dynamic LifeAct::GFP signal in 3° sister dendrites**. Images of PVD::mCherry (left) and PVD::LifeAct::GFP (right) and were collected at 1.3 min intervals with a 100 X objective in a Nikon spinning disk microscope (see Materials and Methods). The length of the video is 60 minutes. The movie was edited using NIS elements with automatic deconvolution. Arrows track 3º dendrites undergoing contact and retraction. Scale bar is 5 µm.

**Movie S5: Time-Lapse video demonstrating the trafficking of UNC-34::mCherry puncta in PVD 3° dendrites.** Images of PVD::mCherry::UNC-34 were captured at 2 minute intervals in a single Z plane with a 100X objective in a iSIM microscope for 3 minutes (See Materials and Methods). Tips of 3° dendrites are indicated by a magenta arrowhead. The movie was edited using a Metamorph and deconvolved using Fiji software. Scale bar is 5 µm.

