## Supplemental Figure 1 for "Actin assembly and non-muscle myosin activity drive dendrite retraction in an UNC-6/Netrin dependent self-avoidance response"

### Figure S1: PVD 3° dendrites undergo self-avoidance in an UNC-6/Netrin dependent manner

A. wild-type

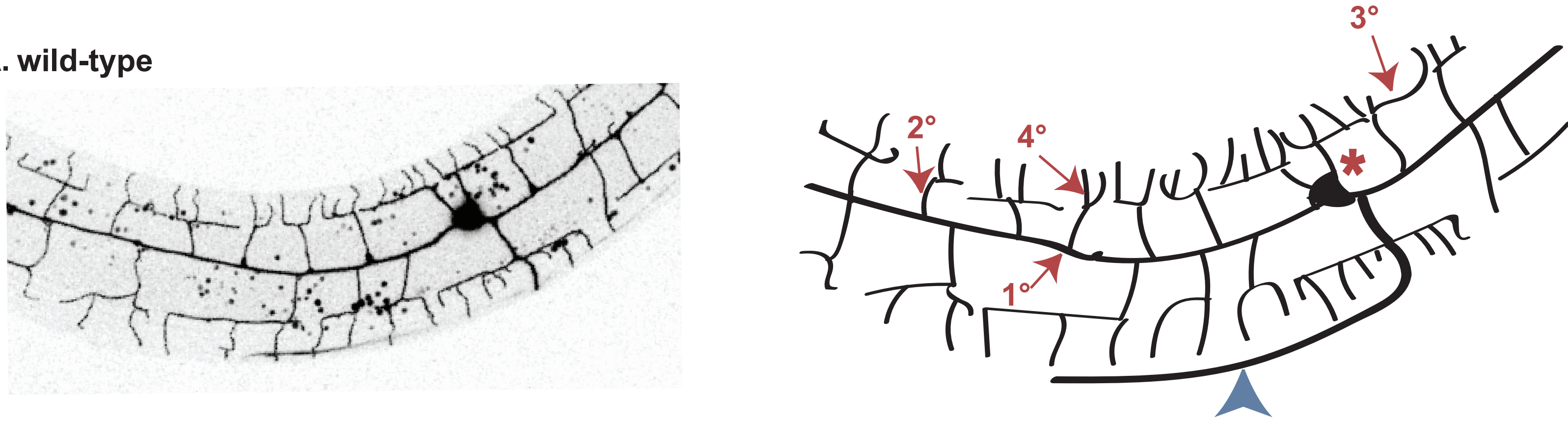


B. *unc-6 (ev400)*

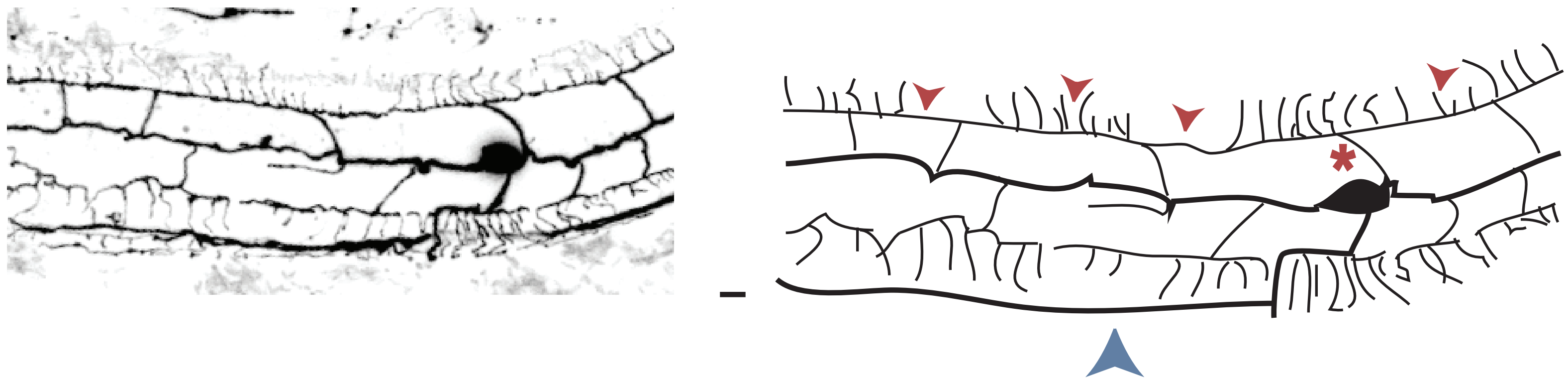


C.

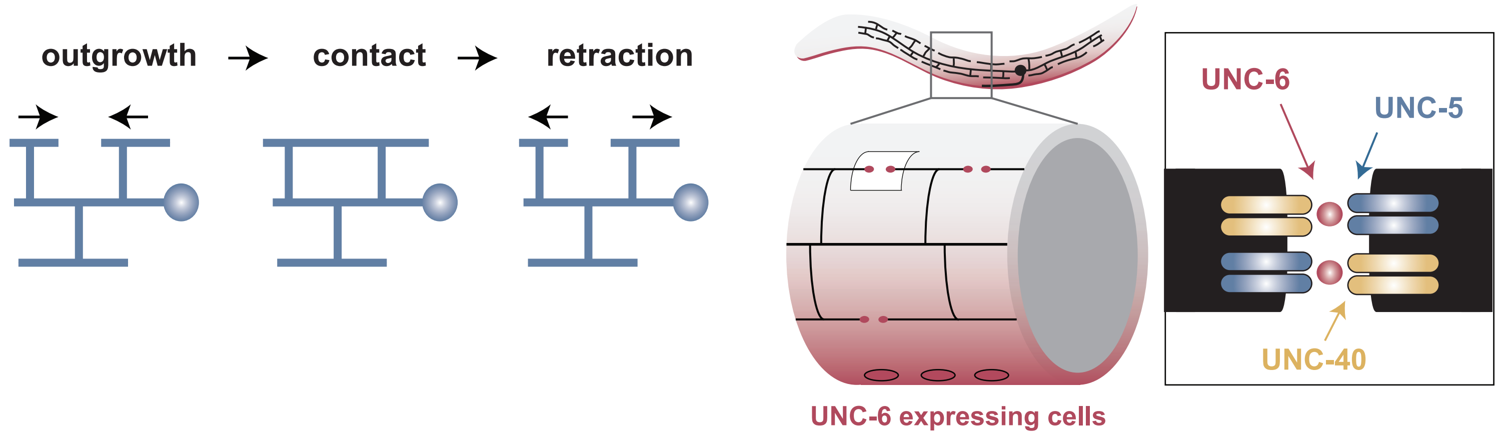
