## Supplementary figures and images for "Actin assembly and non-muscle myosin activity drive dendrite retraction in an UNC-6/Netrin dependent self-avoidance response"

### Supplemental Figure 2

#### Figure S2: Mutations in *unc-34* and *unc-73* disrupt PVD dendrite morphology

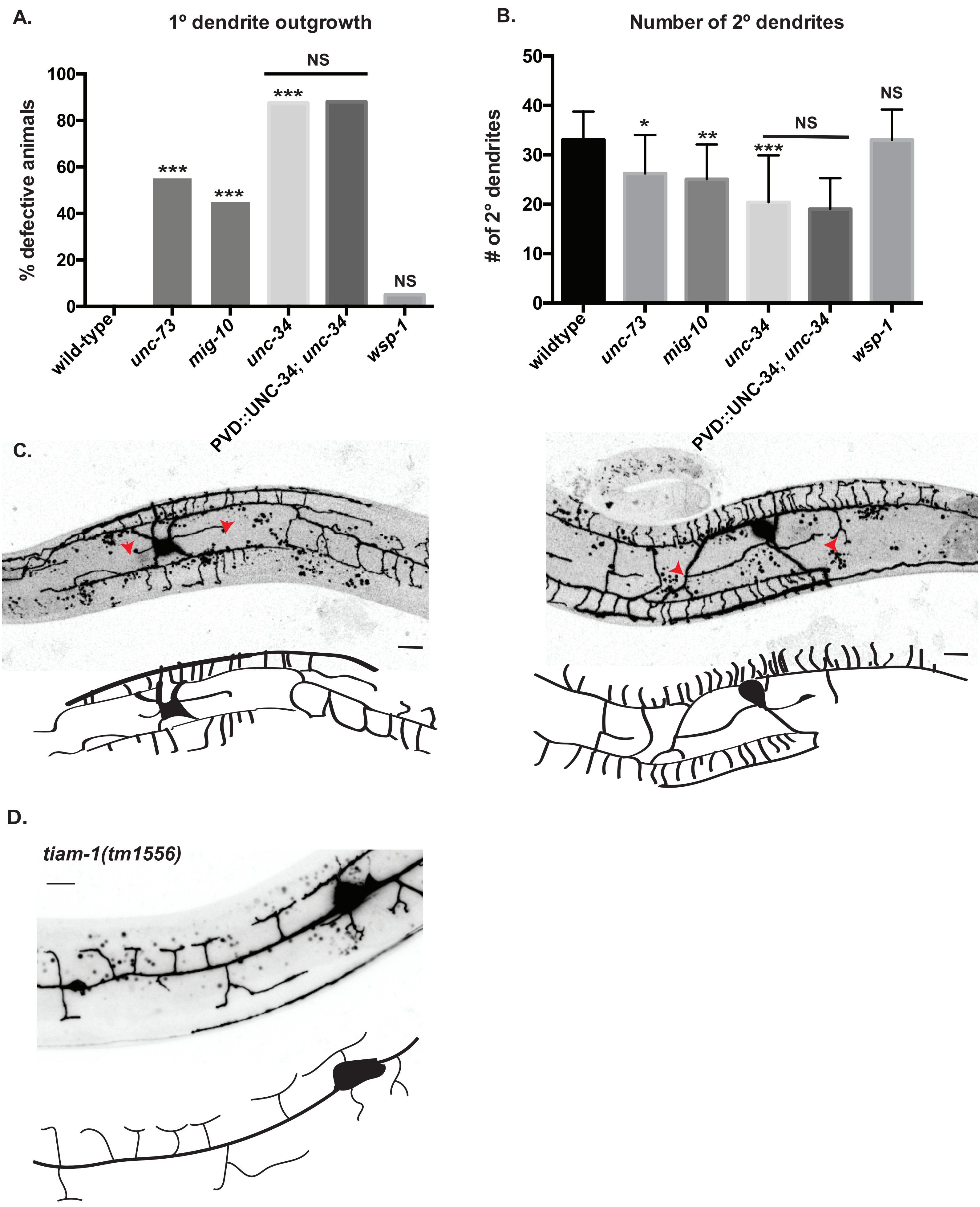

### Supplemental Figure 4

#### Figure S3: Actin dynamics during dendrite self-avoidance

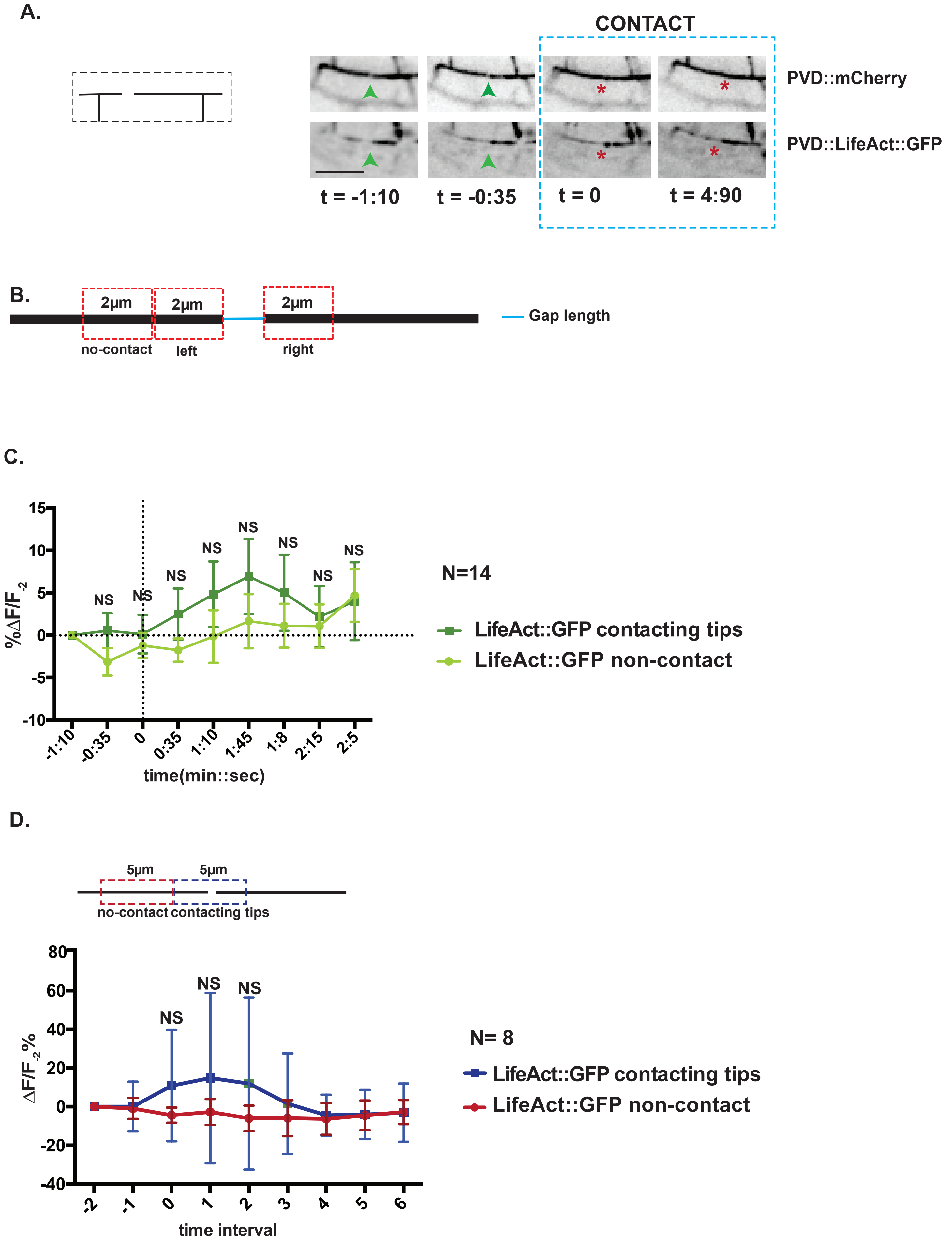

### Supplemental Figure 5

#### Figure S4: Actin dynamics during dendrite self-avoidance.

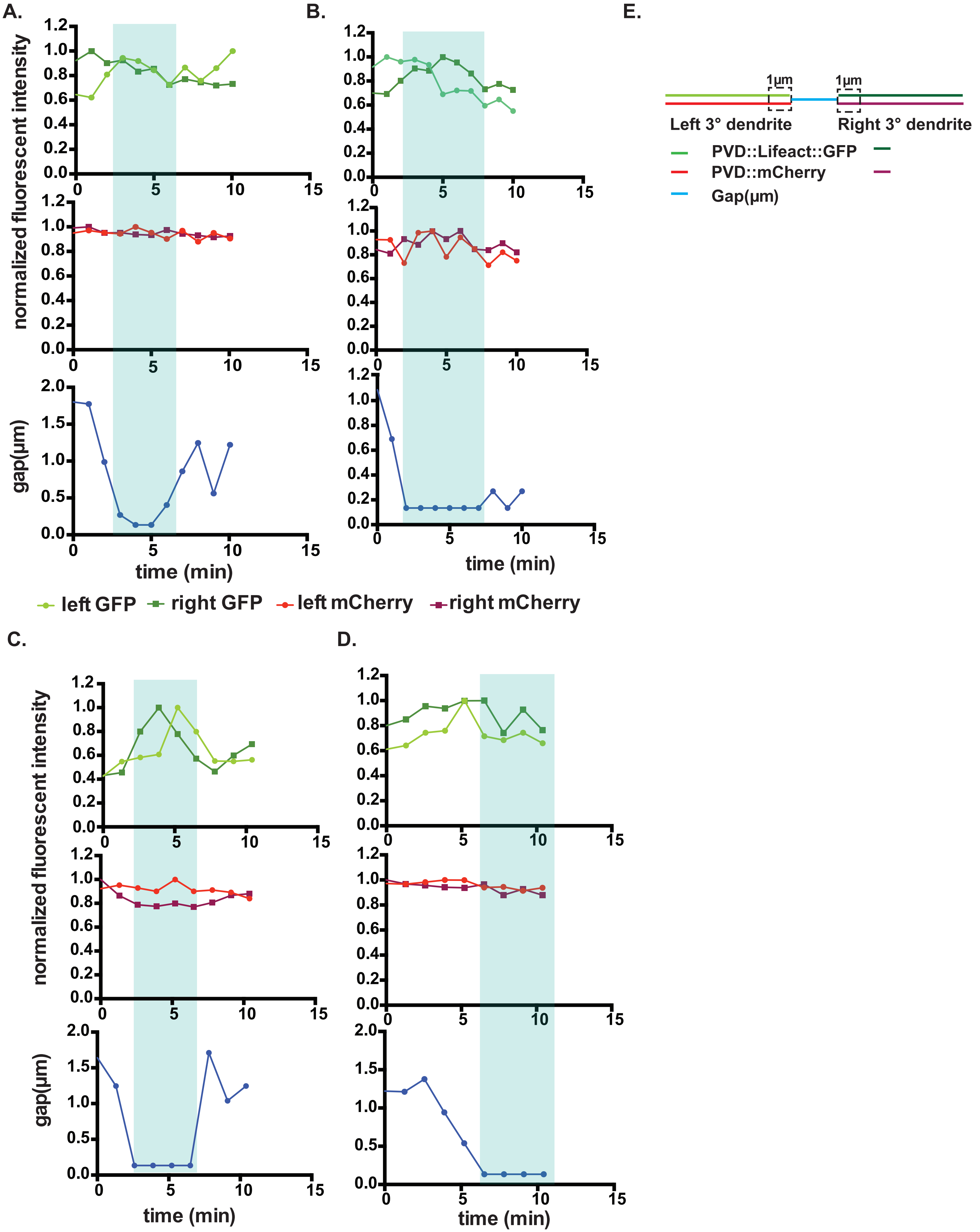
