## Supplemental Figure 3 for "Actin assembly and non-muscle myosin activity drive dendrite retraction in an UNC-6/Netrin dependent self-avoidance response"

**Figure S3: Localization and trafficking of PVD::mCherry::UNC-34 puncta in PVD using Airy Scan microscopy.**

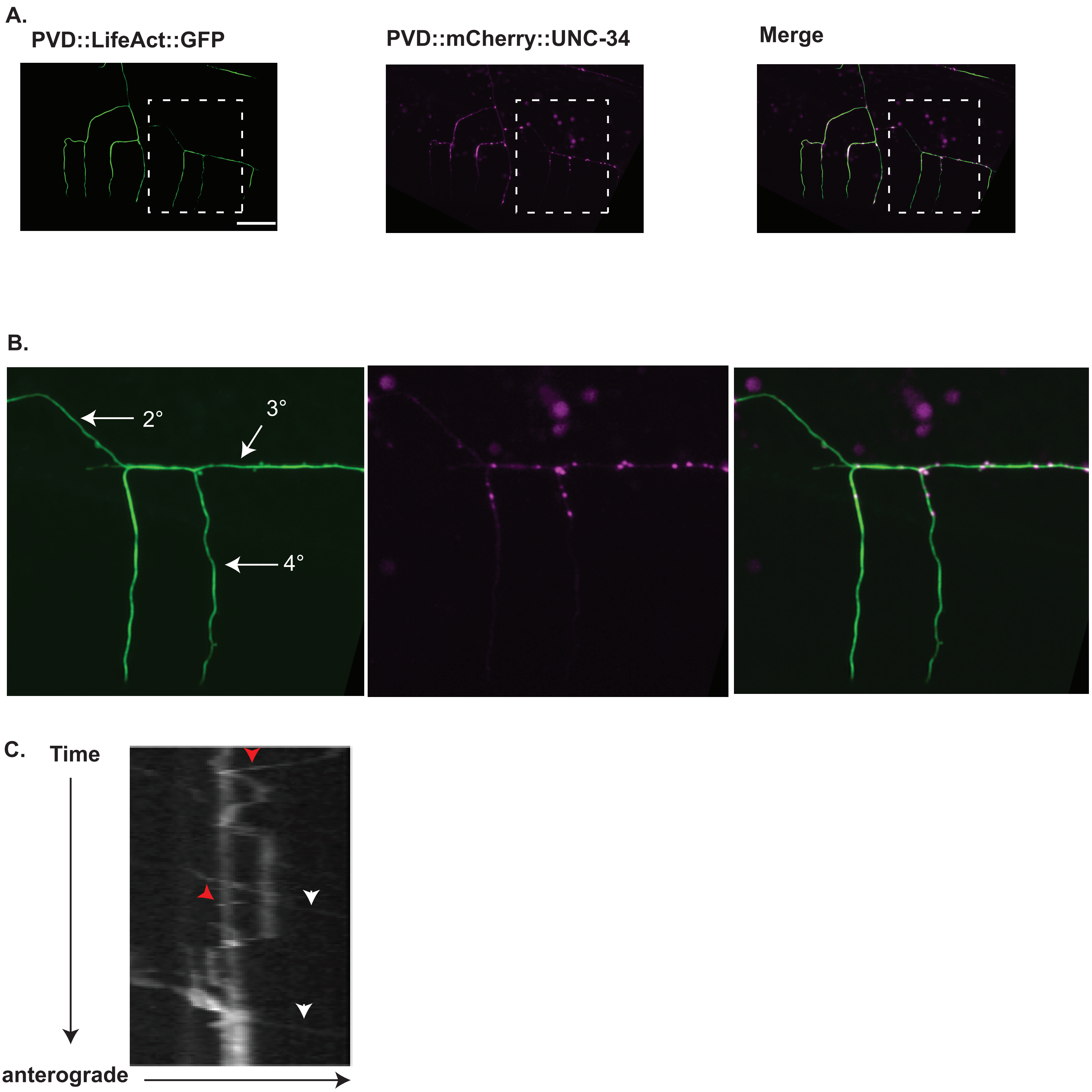
